# A cell-free bacterial signal orchestrates trans-kingdom fitness tradeoff to enhance sulfur deficiency tolerance in plants

**DOI:** 10.1101/2025.05.06.652065

**Authors:** Arijit Mukherjee, Mrinmoy Mazumder, Arun Verma, Hitesh Tikariha, Raktim Bhattacharya, Ooi Qi En, Sanjay Swarup

## Abstract

Plant-associated microorganisms interact with each other and host plants via intricate chemical signals, offering multiple benefits, including plant nutrition. We report such a mechanism through which the rhizosphere microbiome improves plant growth under sulfur (S) deficiency. Disruption of plant S homeostasis resulted in a coordinated shift in composition and S-metabolism of the rhizosphere microbiome. Using this insight, we developed an 18-membered synthetic rhizosphere bacterial community (SynCom) that rescued growth of Arabidopsis and a leafy Brassicaceae vegetable under S-deficiency. This beneficial trait is taxonomically widespread among SynCom members, with bacterial pairs predominantly providing synergistic benefits to host plants. Notably, stronger competitive interactions among SynCom members conferred greater fitness benefits to the host, suggesting a trans-kingdom fitness tradeoff. Finally, guided chemical screening, deletion knockout mutant, and targeted metabolomics identified and validated microbially released glutathione (GSH) as the necessary bioactive signal that orchestrates plant-microbe (trans-kingdom) fitness tradeoff and improves plant growth under sulfur limitation.

**Highlights:**

- Alteration in plant sulfur (S) homeostasis leads to coordinated shifts in the composition and S-metabolism of the rhizosphere microbiome.
- Taxonomically divergent bacterial members within the rhizosphere microbiome improve plant growth under S-deficiency.
- Stronger competitive interactions within rhizosphere bacterial members provide greater fitness benefit to the host plant under S-deficiency, suggesting a trans-kingdom fitness tradeoff.
- Microbially released glutathione (GSH) is a necessary bioactive signal that improves plant growth under sulfur limitation and orchestrates plant-microbe (trans-kingdom) fitness tradeoff.

## Introduction

The rhizosphere, a 2-10 mm region surrounding the root surface, contains a rich repertoire of chemical cues emanating from both plant roots and the colonizing microorganisms. These microorganisms, collectively known as the ‘rhizosphere microbiome’ can influence plant health via microbe-microbe and microbe-plant chemical dialogue. Plant-derived chemicals in both soluble and volatile forms have been shown to influence the assembly of the rhizosphere microbiome^1,2,3,4,5,6,7,8^ with consequent fitness benefit to the host plant.^5,7,8^ The extraordinarily diverse rhizosphere microbiome also produces an array of chemicals that mediate inter-species interactions^9,10^ and influence plant’s responses towards environmental cues.^11,12^

Sulfur (S) as the fourth major macronutrient (after nitrogen, phosphorus, and potassium), plays a pivotal role in plant growth, development, and for agriculturally relevant plant traits.^13^ It is abundant in soil (fifth most abundant element on the Earth^14^), but its plant-available form sulfate, is scarce (comprising only 5% of the total S pool in soil).^15^ This is exacerbated by a recent shift in the global S cycle, leading to widespread S-deficiency in agricultural soils.^16,17^ Plants have evolved physiological, molecular, and metabolic responses to adapt to S-deficient environments. Typical examples include lateral root formation,^18,19^ upregulation of sulfate transporters,^20^ and redistribution of S-metabolites.^21,22^ Overaccumulation of O-acetyl serine (OAS) is a common metabolic response towards S-deficiency in Arabidopsis seeds, shoots, and roots.^23^ In germinating seeds, S is redistributed from S-rich seed storage proteins (SSPs) towards S-poor SSPs.^24^ In both root and shoot, SLIM1, a master transcriptional regulator of S-deficiency responses upregulates Sulfur Deficiency Induced (SDI) proteins to repress glucosinolate biosynthesis.^25,26^ SLIM1, along with the OAS responsive gene *MSA1* increases sulfate uptake in roots, through upregulation of SULTR1;1 and SULTR1;2 transporters.^22^ In addition to these molecular regulators, S provisioning by microbial mineralization of organic-S and bacterial volatile dimethyl disulfide (DMDS) also contribute to alleviation of S-deficiency.^27^ However, the role of the rhizosphere microbiome, chemical signals in nutritional stress and their interactions for adaptation of plants remains poorly understood.

In this study, we investigated the interplay between plant S nutrition and the rhizosphere microbiome. First, using known plant S nutritional mutants and rhizosphere metagenomics, we demonstrated that S-metabolism of the rhizosphere microbiome is coordinated in response to alteration in plant S levels. Building on this, we assembled a functionally representative but reduced 18-membered synthetic community (SynCom) and studied its effects on the growth of Arabidopsis and a leafy Brassicaceae vegetable- *Brassica rapa* var. *parachinensis* (ChoySum) under S-deficiency. By employing the SynCom alongside plate-based gnotobiotic systems, we studied the outcomes of individual SynCom members and selected bacterial pairs on plant growth under S-deficiency. Recognizing the complexity of bacterial interactions even within an 18-member SynCom, we utilized genome-scale metabolic models and machine learning to investigate the role of the emergent properties of their pairwise interactions on plant growth under S-deficiency. Lastly, we combined chemical supplementation, bacterial gene deletion, and targeted metabolomics to identify and validate the role of a bacterial exo-metabolite in promoting plant growth under S deficiency. Our results demonstrate a trans-kingdom fitness tradeoff between plants and bacteria, mediated by the ubiquitous sulfur-containing metabolite glutathione, which acts as a bacterial signal that mitigates the sulfur deficiency response in plants.

## Results

### Arabidopsis genes involved in sulfur nutrition govern the microbiome composition in the root, shoot, and rhizosphere

To investigate the influence of plant S nutrition on bacterial communities in plant niches (root, shoot, and rhizosphere), we selected a literature-curated set of Arabidopsis genotypes affected in S uptake, transport and redistribution, and regulation (epigenetic and transcriptional) of S homeostasis (Figure 1A and Table S1). Relationships between plant sulfate content, microbiome composition and functions across the three niches (root, shoot, and rhizosphere) were then determined in a pot-study of plants grown in commercial black peatmoss soil (see Materials and Methods; Figure 1B). Sulfate levels in these genotypes varied from 11 to 53.12 µmol per mg dry biomass, with a moderate but statistically significant correlation with shoot biomass (Spearman’s l1 = 0.23; p < 0.05*, Figures 1C, S1C, and S1D), suggesting that sulfate levels influence aboveground biomass. As anticipated, genotypes within the same functional group exhibited similar sulfate accumulation patterns (Figure 1C). Particularly, *msa1-3* and *sdi2;1* showed significant over-accumulation of sulfate in shoot (Cohen’s ES ≥ 0.5; q ≤ 0.1), while *sultr2;1* and *sultr2;2* accumulated less sulfate (Cohen’s ES ≤ 0.5; q < 0.1) than the *wild-type* Col-0 plants (Figure 1D). Notably, none of these genotypes accumulated significantly different nitrate or phosphate levels than the Col-0 plants (Figure 1D; Figures S1A and S1B). Consequently, we selected five genotypes (*msa1-3*, *sdi2;1*, *sultr2;1*, *sultr2;2*, and Col-0) for microbiome profiling across the root, shoot, and rhizosphere (Figure 1B).

**Figure 1.**
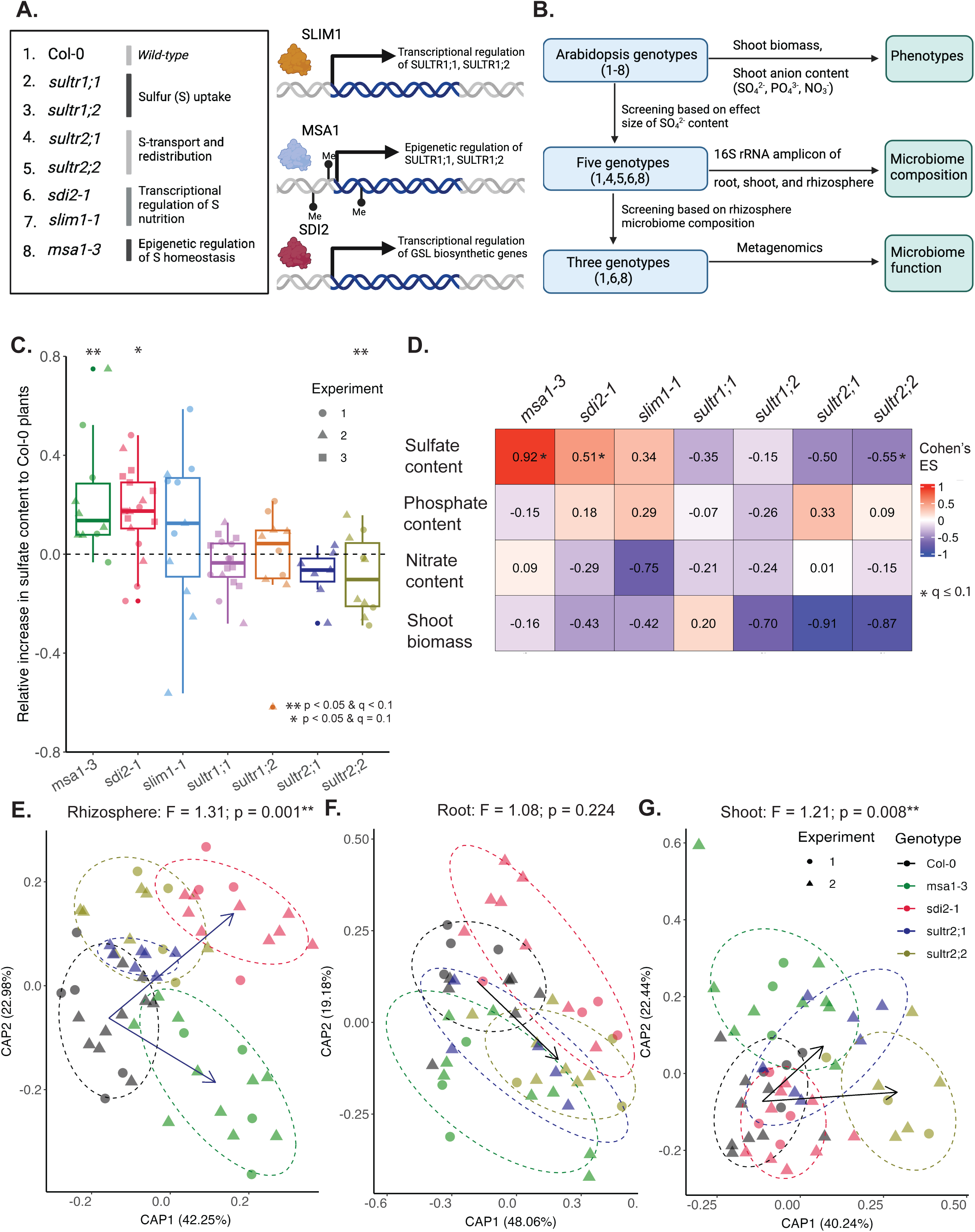
Arabidopsis genes involved in sulfur nutrition govern the microbial community composition in the root, shoot, and rhizosphere. **A.** Arabidopsis genotypes involved in different processes of plant S nutrition, tested in this study. Genotypes are grouped according to their role in S nutrition processes. **B.** Workflow for screening Arabidopsis genotypes to identify relationships between plant S nutrition, microbiome composition, and functions. Genotypes were first screened based on their shoot sulfate content and five genotypes were used for their microbiome composition analysis in three plant niches. Based on this, the rhizosphere microbiome of three genotypes were subjected to metagenomics analysis to infer their potential functions. **C.** Boxplot showing shoot sulfate content in all genotypes. Statistical significance (denoted by letters on top of the boxes) was determined using ANOVA, controlling for experiments, followed by a post-hoc Tukey’s test (q ≤ 0.1). Data points are colored according to genotypes with the following number of biological replicates: Col-0 (n = 18), *msa1-3* (n = 10), *sdi2;1* (n = 18), *slim1-1* (n = 11), *sultr1;1* (n = 16), *sultr1;2* (n = 8), *sultr2;1* (n = 7), and *sultr2;2* (n = 10), distributed across two independent experiments (indicated by the shape of data points). Three genotypes-Col-0, *sdi2;1* and *sultr1;1* were included for three independent experiments. Plants were grown in commercial Black peatmoss soil for five weeks at 21°C (16 h light / 8 h dark cycle) before sampling. **D.** Matrix showing the Cohen’s effect size (ES) of anion content (sulfate, phosphate, and nitrate) and shoot biomass for all genotypes compared to wild-type Col-0 plants. Cells are colored according to ES and asterisk indicates statistical significance (q ≤ 0.1). Detailed data for sulfate, phosphate, nitrate levels, and shoot biomass are shown in Figures 1C, S1A, S1B, and S1C, respectively. **E,F,G**. Constrained ordination of microbiome composition for selected genotypes in the rhizosphere (E), root (F), and shoot (G). Black arrows connect the centroids of the most separated microbiomes along CAP1 and CAP2 ordination axes. Data points are colored according to the genotypes and ellipses show the parametric smallest area around the mean that contains 80% of probability mass for each genotype. Statistical significance (F-values) was calculated using PERMANOVA and are shown at the top of the plots. Number of biological replicates for rhizosphere (E), root (F), and shoot (G), respectively: Col-0 (n = 15, 11, and 13), *msa1-3* (n = 14, 11, and 11), *sdi2;1* (n = 13, 12, and 13), *sultr2;1* (n = 7, 5, and 7), and *sultr2;2* (n = 11, 10, and 8). Data are shown for two independent experiments indicated by shape of the data points.

Consistent with earlier findings, alpha diversity of bacterial communities in plant niches were significantly lower than that of the bulk soil microbiome (Figures S1F, S1G, and S1H for root, shoot, and rhizosphere, respectively).^28,29,30^ However, no significant differences in alpha diversity (Shannon index) were observed among the genotypes across these niches (Figures S1F, S1G, and S1H). Beta-diversity analyses using constrained ordination, however, showed that the composition of bacterial communities in the rhizosphere, but not other niches, was affected by *sdi2;1* (F = 1.87, p = 0.001**) and *msa1-3* (F = 1.49, p = 0.012*) (Figure 1E and Table 1). This suggests that the loss of transcriptional and epigenetic regulators of S homeostasis, *sdi2;1* and *msa1-3* respectively had significant effects on the bacterial communities in a niche-specific manner. Conversely, processes related to S transport and redistribution significantly impacted the microbial community composition in the shoot (*sultr2;1*: F = 1.55, p = 0.02* and *sultr2;2*: F = 1.71, p = 0.001**; Figure 1F and Table 1) and root (*sultr2;2*: F = 2.18, p = 0.009**; Figure 1G and Table 1). Interestingly, variations in microbiome composition across all niches were better explained by plant genotypes, rather than accumulation of sulfate in shoot (Table S3), suggesting that genes involved in sulfur nutrition significantly influence the composition of bacterial communities in these niches. The microbiome composition of these genotypes was better separated in the rhizosphere (F = 1.31, p = 0.001***; Figure 1E) than other niches (root: F = 1.08, p = 0.224; Figure 1F, shoot: F = 1.21, p = 0.008**; Figure 1G), therefore, we focused our analyses on the rhizosphere microbiome in subsequent studies. Overall, our results demonstrate that Arabidopsis genes associated with S nutrition significantly influence microbiome (bacterial) composition across the niches.

**Table 1.** Results from pairwise PERMANOVA for microbiome composition of all genotypes compared to wild-type Col-0 plants across rhizosphere, root, and shoot.

| Comparisons | Rhizosphere | Root | Shoot |
| --- | --- | --- | --- |
| Col-0 vs *msa1-3* | F = 1.49; p = 0.012** | F = 0.80; p = 0.695 | F = 0.98; p = 0.609 |
| Col-0 vs *sdi2;1* | F = 1.87; p = 0.001*** | F = 1.52; p = 0.06 | F = 1.03; p = 0.221 |
| Col-0 vs *sultr2;1* | F = 0.91; p = 0.7 | F=0.74;p=0.829 | F = 1.55; p = 0.035* |
| Col-0 vs *sultr2;2* | F = 1.08; p = 0.28 | F = 2.18; p = 0.009** | F = 1.71; p = 0.001*** |

### Over-accumulation of sulfate in plants coincides with a shift in sulfur metabolism of the rhizosphere microbiome

Given the markedly different microbiome composition was associated with over-accumulation of sulfate (in *msa1-3* and *sdi2;1*), we conducted metagenomic analyses to investigate functional changes in their rhizosphere microbiome. Constrained ordination based on predicted Clusters of Orthologous Genes (COGs) from the metagenome analyses revealed a significant difference in overall functions of the *msa1-3* and *sdi2;1* rhizosphere microbiome (F = 1.25, p = 0.05*; Figure 2A). Among the 1017 differentially represented COGs (metagenomeSeq: q < 0.1 combined from *msa1-3* vs Col-0 and *sdi2;1* vs Col-0; Table S4), 20 GO categories were represented (Table S5). Out of these, translational processes, transport and metabolism of amino acids, lipids, and nucleotides were the significant ones (Fisher’s test: q < 0.1; Figure 2B; Table S5). The category of ‘’transport and metabolism of amino acids’’ was the top one among the metabolism-related processes which represented 10.06% of differentially represented COGs (q = 0.0007; Figure 2B and Table S5). To investigate these amino acid subclasses further, we focused on the 40 COGs within this category. Among these, cysteine/methionine metabolism-related genes represented 22.50% of the COGs, greater than any other amino acid transport and metabolism related COGs (Figure 2B and Table S6). This highlights the differential representation of major S-metabolism related COGs (Cys/Met metabolism) in the rhizosphere metagenomes of sulfate over-accumulating genotypes. Next, we asked the question in addition to those involved in Cys/Met metabolism, which other S-related genes are differentially represented in *msa1-3* and *sdi2;1* rhizosphere metagenomes. In addition to the Cys/Met metabolic genes, those involved glutathione (GSH) metabolism (Glutathione S transferase or GST) and other S-related genes were differentially represented in these two genotypes (combined from KEGG and COGs, metagenomeSeq: q < 0.1; Table S7), highlighting their potential effects on the bacterial S-metabolism in the rhizosphere (Figure 2C). Overall, our findings suggest a cross-kingdom interplay of S-metabolism in the rhizosphere. However, it is important to consider that the observed shifts in rhizosphere microbiome function could result from either direct or indirect effects through pleiotropy in these two genotypes, as MSA1 is an epigenetic regulator of S homeostasis,^20^ whereas SDI2 negatively regulates glucosinolates biosynthesis.^26^

**Figure 2.**
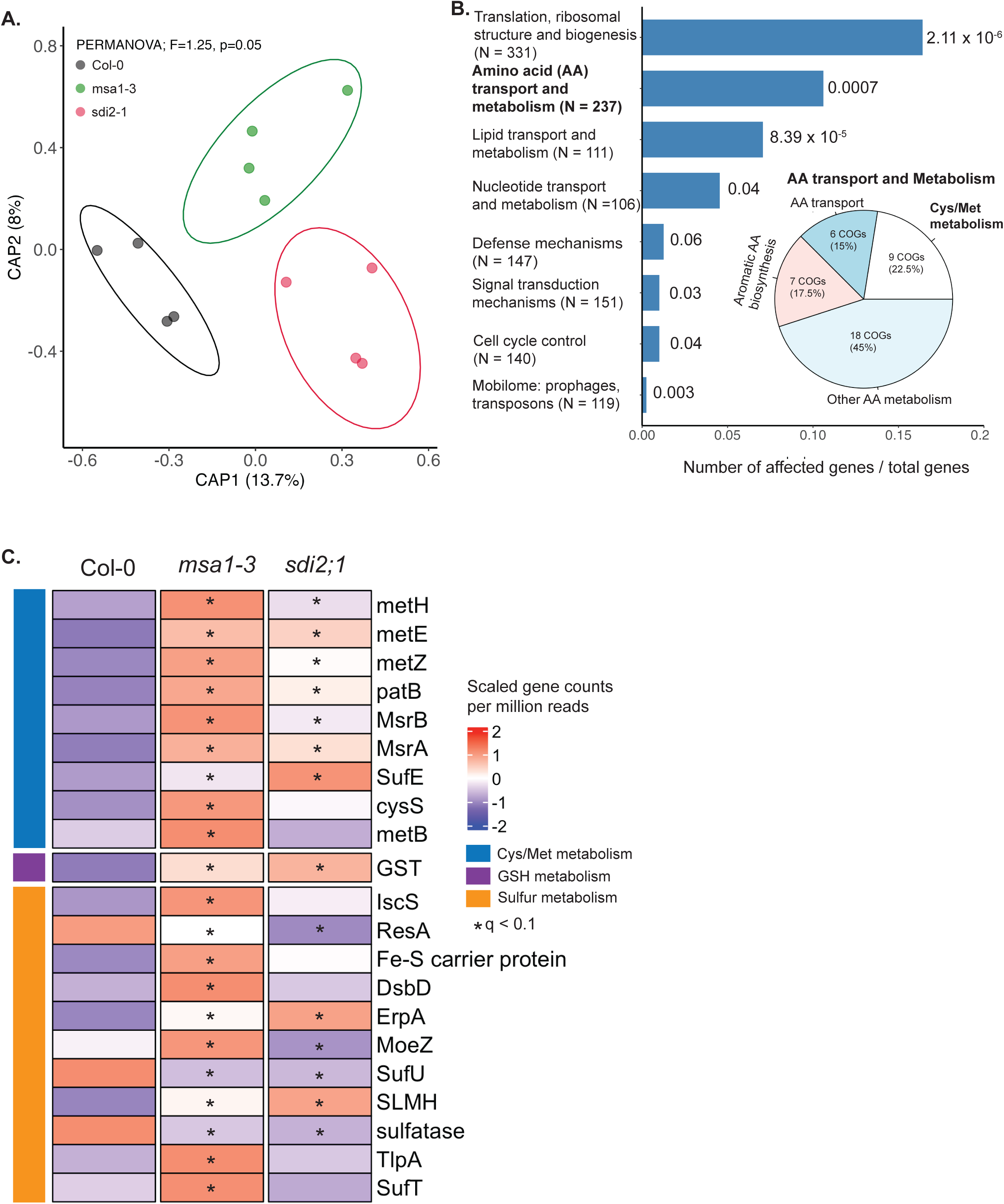
Over-accumulation of sulfate in shoot coincides with a coordinated shift in sulfur metabolism of the rhizosphere bacterial community. **A.** Constrained ordination plot (constrained analysis of principal coordinates or CAP) based on predicted cluster of orthologous genes (COGs) in the rhizosphere metagenomes of Col-0 and the high-sulfate accumulating *msa1-3* and *sdi2;1* (n = 4 each) plants. Data points are colored according to genotypes and ellipses show the parametric smallest area around the mean that contains 80% of probability mass of each genotype. Statistical significance was calculated using PERMANOVA (F = 1.25; p = 0.05*). **B.** Bar plot showing the over-representation of functional categories of COGs (y-axis) in *msa1-3* and *sdi2;1* rhizosphere metagenomes, compared to Col-0 plants. The proportion of COGs over-represented in each category is shown along the x-axis. Statistical significance was calculated based on Fisher’s exact test (q < 0.1) and are indicated beside the bars. The total number of COGs within each category is shown in brackets. The pie chart in the inset shows the distribution of differentially represented COGs within the amino acid (AA) transport and metabolism category. The number of COGs and percentage distribution within each sub-category are shown, with the largest category Cys/Met metabolism shown in bold face. **C.** Heatmap showing the differential representation of S-metabolic genes in the rhizosphere metagenomes of Col-0, *msa1-3,* and *sdi2;1* plants. Colors within the heatmap represent scaled mean counts of differentially represented genes in either *sdi2;1* or *msa1-3*, or both compared to Col-0 plants (q < 0.1*). The differentially represented genes are grouped according to the three S-metabolic processes.

### A functionally representative, but reduced synthetic rhizosphere bacterial community rescues plant growth under sulfur-deficiency

As the host sulfate levels affected the microbiome composition and S-metabolism, we asked whether active manipulation of the rhizosphere microbiome through a synthetic community, representative of S-metabolic genes, could provide feedback to the plants under different sulfur levels. We assembled an 18-membered synthetic rhizosphere bacterial community (SynCom) named <u>S</u>ulfur <u>P</u>roteobacteria <u>A</u>ctinobacteria <u>F</u>irmicutes 18 (SPAF18) that represented 80% of the S-metabolic genes from a global-scale collection of rhizosphere microbiome^9^ (see Materials and Methods; Figures S3C and S3D) and the diversity of S-metabolic genes present in the high and medium-quality metagenome-assembled genomes (MAGs) (Figures S2 and S3E) from the pot-study with the selected genotypes described above (Figure 2).

Under S-deficiency in a plate-based gnotobiotic setup, a significant reduction in shoot area and fresh biomass in Arabidopsis were observed compared to S-sufficient conditions (Figures 3B, 3C, and 3D). However, root phenotypes-root network, primary root length, and number of lateral roots remained unaffected (Figures S4A, S4B, and S4C). These findings align with the previously reported S-deficiency phenotypes in Arabidopsis^19^, validating the suitability of this plate-based gnotobiotic setup for studying plant-microbe interactions under S-deficient conditions.

**Figure 3.**
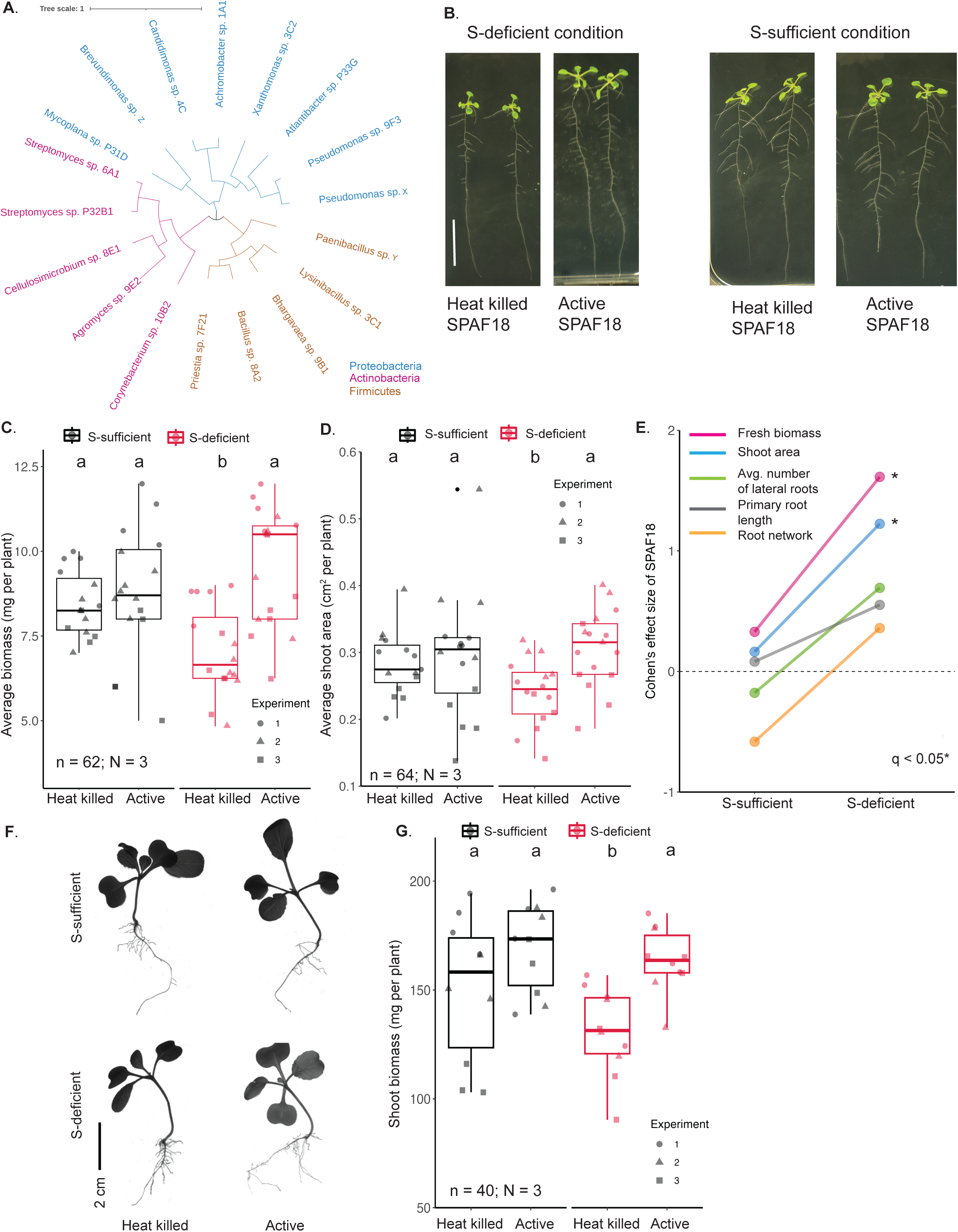
A functionally representative but reduced synthetic rhizosphere bacterial community rescues plant growth, specifically under sulfur-deficiency. **A.** Phylogenetic tree of 18 members of the SPAF18 SynCom. Branch colors indicate their taxonomic affiliations to the three phyla. **B.** Image of representative Arabidopsis plants (12 days old) inoculated with either heat-killed or active SPAF18 grown under both S-sufficient and S-deficient conditions. **C**. Average biomass of plants (in mg per plant) inoculated with either heat-killed or active SPAF18 in S-sufficient and S-deficient conditions. Number of biological replicates, n = 63 distributed across three independent experiments, N=3 (indicated by shape of data points). Statistical significance (denoted by letters at top of the boxes) was determined using ANOVA, controlling for experiments, followed by a post-hoc Tukey’s test (q < 0.05). **D.** Shoot area (in cm^2^) for individual plants inoculated with either heat-killed or active SPAF18 in S-sufficient and S-deficient conditions. Number of biological replicates, n=64 across three independent experiments, N = 3 (indicated by shape of data points). Statistical significance was determined the same as for C and is indicated with letters on top of the boxes. **E.** Cohen’s effect size of SPAF18 for five root and shoot parameters (shown by different colored lines) measured under both S-sufficient and S-deficient conditions. Statistical significance relative heat-killed SPAF18 is indicated with an asterisk (q < 0.05). **F.** Image of representative ChoySum plants (14 days old) inoculated with either heat-killed or active SPAF18 in both S-sufficient and S-deficient conditions. **G.** Average biomass of ChoySum plants (mg per plant) inoculated with either heat-killed or active SPAF18 in S-sufficient and S-deficient conditions. Number of biological replicates, n = 40, distributed across three independent experiments, N = 3 (indicated by shape of data points). Statistical significance was determined the same as for C and is indicated with letters at top of the boxes.

Remarkably, one-week after co-culturing Arabidopsis plants with SPAF18, the plants appeared healthier with larger rosettes than those treated with heat-killed SPAF18 controls, under S-deficient condition (Figure 3B). However, no differences in plant phenotypes were observed between heat-killed and active SPAF18 treatments under S-sufficient conditions (Figure 3B). These suggest that SPAF18 shifts from a commensal to beneficial role, under S-deficiency. SPAF18 treatment also led to a higher fresh biomass and shoot area under S-deficient conditions (Cohen’s ES = 1.61, q < 0.05; Cohen’s ES = 1.22, q < 0.05, respectively) but not under S-sufficient conditions (Figures 3C, 3D and Table S8). Root traits such as primary root length, root network, and the number of lateral roots also increased, with small to moderate effect sizes (Cohen’s ES = 0.55, 0.35, and 0.69, respectively) though these changes were not statistically significant (Figure 3E, Figures S4A-C, and Table S8). This indicates that the effects of SPAF18 are more pronounced in shoot than in the roots (Figure 3E) and both biomass and shoot area explain most of the variations in the effects of SPAF18 on Arabidopsis phenotypes (Figure S4D).

To investigate whether SPAF18 could rescue plant growth in another Brassicaceae family member ChoySum, we compared the shoot biomass of treated vs heat killed controls. As in the case of Arabidopsis, the shoot phenotypes of the SPAF18 treated plants showed better growth than the controls for S-deficient conditions, but not S-sufficient conditions (Figure 3F). The shoot biomass was significantly higher in the SPAF18 treated plants than the controls for the S-deficient conditions (Figure 3G). Hence SPAF18 has both genomic capacity of S-metabolism and phenotypic capacity to improve S-deficiency tolerance in the host plants.

### Synergistic improvements in plant biomass under sulfur deficiency are most prevalent among SPAF18 member pairs

We assessed the role of individual SPAF18 members in influencing five different plant phenotypes under S-deficiency. Of the 18 strains tested, 13 significantly (q < 0.05) improved at least one of the five phenotypes. Most strains enhanced fresh biomass and shoot area, mirroring the overall effects of SPAF18 (Figure 4A and Figures S5A-E). All Proteobacteria strains significantly increased fresh biomass (Figures 4A and S5A). In contrast, Actinobacteria and Firmicutes included both growth-promoting strains and those without this capacity (Figure 4A and Figures S5A-E). These suggest that the ability to improve plant growth under S-deficiency is taxonomically widespread and functionally redundant among SPAF18 members, as evidenced by their negligible phylogenetic signals (Pagel’s λ = 10^-5^ to 10^-4^ for all five phenotypes). Importantly, majority of these strains failed to improve fresh biomass under S-sufficient conditions, reflecting their similar beneficial capacities within a full community (Figure S5F). Notably, the growth rates of individual strains (in S-deficient medium supplemented with Arabidopsis root extracts) did not correlate significantly with any of the plant phenotypes (Spearman’s ρ = 0.38, 0.24, −0.10, 0.02, and 0.44 for fresh biomass, shoot area, primary root length, number of lateral roots, and root network, respectively; p > 0.05). Therefore, the capacity of SPAF18 members to grow in this media does not reflect their ability to rescue plant phenotypes under S-deficiency.

**Figure 4.**
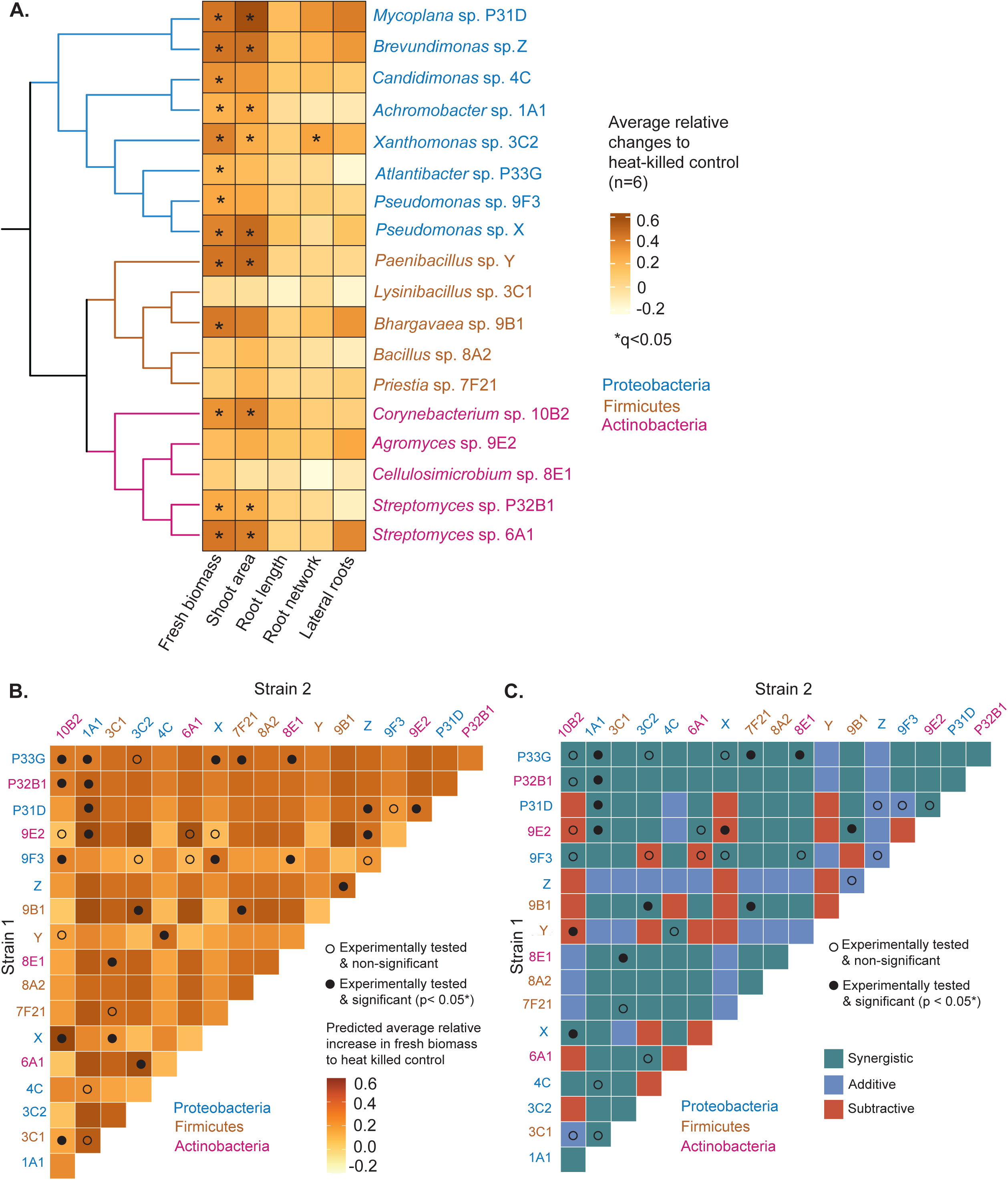
Synergistic improvements in plant biomass under sulfur deficiency are most prevalent among SPAF18 member pairs. **A.** Heatmap showing the effect of individual members of SPAF18 on plant phenotypes compared to heat killed strains under S-deficient condition. Average relative increase (n = 6) to heat killed strains are indicated. Statistical significance was determined using ANOVA followed by a post-hoc Tukey’s test and is indicated with an asterisk (q < 0.05). Strains are arranged based on their phylogenetic affiliation (at the phylum level). **B.** Lower triangle heat map showing the predicted relative increase in fresh biomass for all possible bacterial pairs within SPAF18 under S-deficiency. The predicted values are based on a random forest regression model (Table S8). Experimentally tested pairs and their statistical significance compared to heat killed controls are indicated. **C.** Lower triangle plot showing the effects of bacterial pairs on plant biomass compared to their individual strains. Heatmap cells are colored according to their effects-Synergistic: Increase in plant biomass for bacterial pairs are more than their individual strains; Additive: Increase in plant biomass for bacterial pairs are within the effects of individual strains; Subtractive: Increase in plant biomass for bacterial pairs are less than their individual strains. Experimentally tested pairs and their statistical significance compared to individual strains are indicated.

Given the taxonomically widespread distribution and individual variations of these beneficial effects within SPAF18, we asked whether interactions among them could lead to emergent properties with consequent effect on the host. For this purpose, we tested the effects of representative pairs on plant biomass under S-deficiency (See Materials and Methods). To limit the experimental pairs, a subset was selected by clustering strains based on their performance in mono-association assays and selecting representative strains from both within and between clusters (Figure S7A and Table S9). We then developed machine learning models to predict the effects of the untested pairs (see Materials and Methods). Among the 15 models tested, the random forest regression performed best in terms of its training and testing accuracy (Training and testing R^2^ = 0.77 and 0.44, respectively; Table S9). Albeit, the predictive accuracy was moderate for the untested combinations, as has been found previously^31^, we used this model to predict the outcomes of all pairwise combinations on fresh biomass under S-deficiency (total number of possible pairwise combinations, out of a total of 18 strains i.e., ^18^C_2_ = 153) (Figure 4B and Table S10). The effects of bacterial pairs on fresh biomass ranged from synergistic and additive to subtractive, compared to their individual effects (Figure 4C). Notably, the majority (60.3%) of these pairs provided greater benefit to the plants compared to their individual members, suggesting a prevalence of synergistic benefit to the host plant (Figure 4C). However, we also noted that many of those experimentally verified pairs were not statistically significant. Apart from these synergistic effects, both additive (greater benefit to the plant compared to one of the strains: 21.6%) and subtractive (lower benefit to the plant compared to their individual effects: 17.6%) effects were observed (Figure 4C). This suggests that pairwise interactions among SPAF18 members lead to diverse emergent properties, with synergistic effects on the host being prevalent.

### Competitive interactions within SPAF18 enhance fitness benefit to the host plant under S-deficiency: A trans-kingdom fitness tradeoff model

The above observation that interactions among bacterial pairs give rise to emergent functions, prompted us to investigate the relationship between the nature of their metabolic interactions and their consequent effect on host fitness. To explore this, we first predicted pairwise interaction scores among the members of SPAF18 based on their simulated growth rates using genome-scale metabolic models (GSMMs), which were subsequently validated experimentally (Figure 5A). GSMMs were simulated in a custom medium (MS supplemented with artificial root exudates) designed to replicate our experimental setup (Figure 5A; Tables S11 and S12). To investigate the relationship between bacterial interactions and plant fitness, we experimentally assessed the fitness benefits to host plants under S-deficiency for 19 bacterial pairs, randomly selected from their range of interaction scores (Table S13). Interactions were deemed negative if coculture growth rates were at least 10% lower than in monoculture, and vice versa (See Materials and Methods for details). Nearly all predicted interactions were negative (215 out of 240) with the majority being competitive (81.66%; Figures 5B, 5C, S7B, and Table S12). Most of the remaining interactions were amensal (15.83%), with only a small fraction being neutral (2.50%) (Figure 5B). Experimental growth studies of *Pseudomonas* sp. X in combination with the remaining 15 strains in the same medium as used for the *in-silico* predictions (MS supplemented with artificial root exudates) further validated the predicted outcomes (see Materials and Methods; Table S14 and Figure S7C). Similar validation for amensal combinations could not be tested due to lack of a distinguishing marker of the strains. These interaction scores ranged from - 0.42 to 0.02 (Table S13), allowing us to examine the relationship between the strength of these interactions and their effect on fresh biomass under sulfur deficiency. Notably, the interaction strengths (calculated as the sum of interaction scores for each pair) showed a negative correlation with the relative fitness benefit (calculated as mean relative increase in biomass to heat killed controls) to the host plant (Pearson’s r = - 0.45, p = 0.049*; Figure 5D). This suggests that stronger competitive interactions (more negative sum of interaction scores) among bacterial pairs further enhance plant fitness under S-deficiency. As stronger competition reduces bacterial reproductive fitness, we conclude that host plant fitness improves at the expense of bacterial fitness, highlighting a trans-kingdom fitness tradeoff.

**Figure 5.**
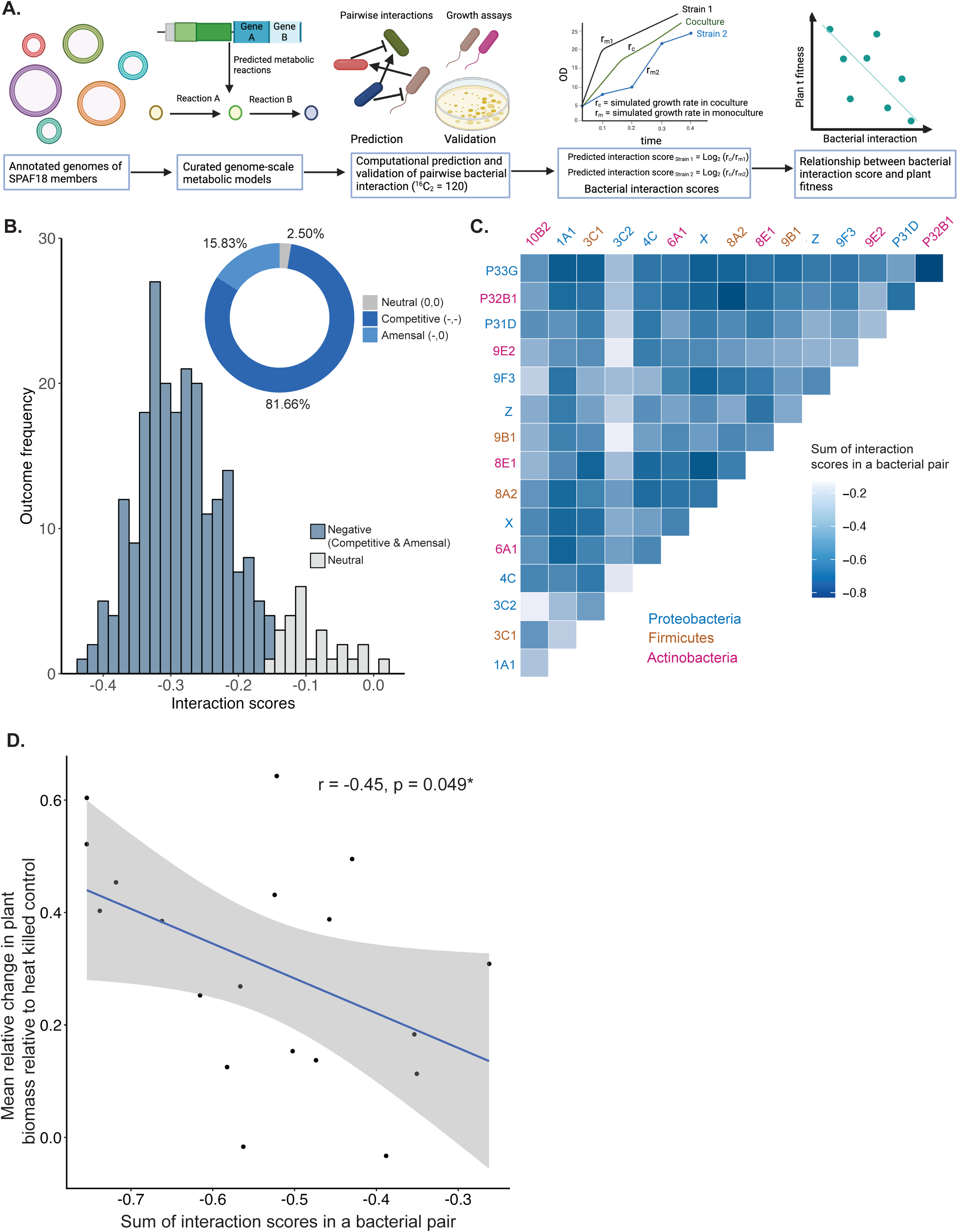
Competitive interactions within SPAF18 enhance fitness benefit to the host plant under S-deficiency: A trans-kingdom fitness tradeoff model. **A.** Schematic showing the approach used to infer the relationship between bacterial interactions and plant growth under S-stress. Genome-sequenced isolates of SPAF18 were used to develop genome-scale metabolic models and computational prediction of their pairwise bacterial interactions based on their simulated growth rates in Artificial root exudates + MS media (Table S12). Selected pairwise interactions were validated using growth assays (Table S14). Interaction score for each strain in a bacterial pair was calculated by calculating Log_2_ of the proportion of their growth rates in coculture and monoculture. These interaction scores were then used to infer their relationship with enhancement in plant fitness. **B.** Distribution of interaction scores among selected bacterial pairs within SPAF18 (^16^C_2_ = 120). Histogram displays the frequency distribution of pairwise interaction scores of the selected members and the donut plot displays the distribution of ecological interactions among the bacterial pairs. **C.** Lower triangle heatmap showing the sum of interaction scores in a bacterial pair. Strains are colored according to their phylogenetic affiliations. **D.** Scatterplot showing the relationship between mean relative increase plant biomass and the sum of interaction scores of bacterial pairs. Pearson’s correlation coefficient (r = −0.45) and statistical significance (p = 0.049*) are indicated.

### Cell-free glutathione from bacterial exudates is a necessary signal that improves plant growth under S-deficiency through plant-microbe fitness tradeoff

SPAF18 exhibited a similar capacity to enhance plant biomass under S-deficiency, whether in direct contact with the host plant or through its cell-free extract (Figure 6A). This suggests that the beneficial effect may be mediated by some released bioactive molecule(s). However, no detectable sulfate was found in the cell-free extract of SPAF18 (Figure S6C) and sulfate levels of Arabidopsis plants didn’t change significantly with treatment of SPAF18 or its selected individual members (S6A and S6B). These indicate that the beneficial effect is not mediated by released sulfate and its concomitant uptake by host plants. Furthermore, the phenotypes of Arabidopsis plants treated with no bacteria and those treated with heat-killed controls were indistinguishable (Figure S6D), ruling out fertilization or Microbe-Associated Molecular Pattern (MAMP)-mediated effects.

**Figure 6.**
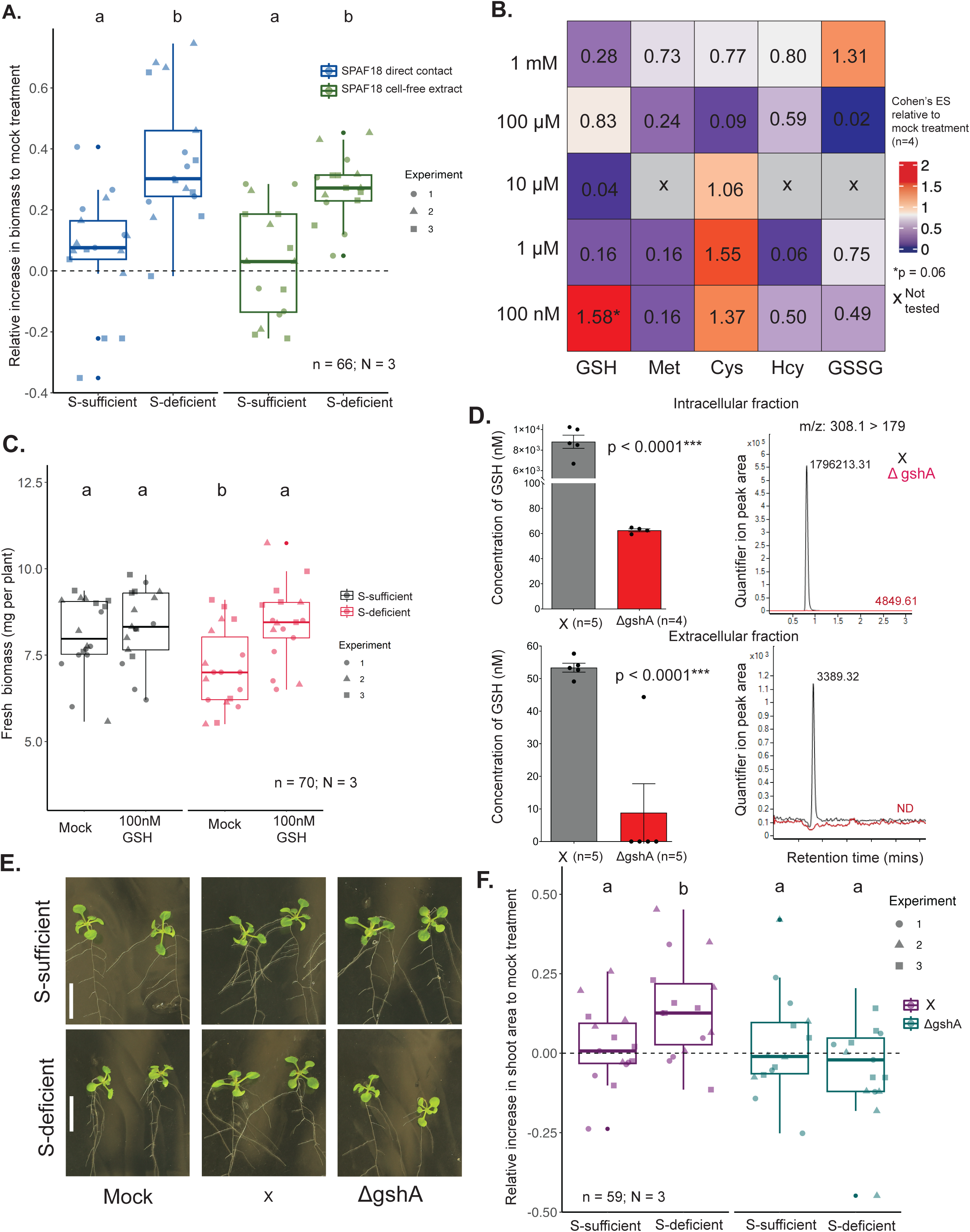
Diffusible glutathione from bacterial exudates is a necessary signal that stimulates plant growth under S-deficiency. **A.** Boxplot showing the relative growth rescue by SPAF18 either in direct contact (blue boxes and data points) or through cell-free extract (green boxes and data points) in both S-sufficient and S-deficient conditions. Number of biological replicates, n = 66 across three independent experiments, N = 3 (indicated by shape of data points). Statistical significance (denoted by letters on top of the boxes) was determined using ANOVA, controlling for experiments, followed by a post-hoc Tukey’s test (q < 0.05). **B.** Heatmap showing the Cohen’s effect size (ES) for the tested S-metabolites at different concentrations (nM to mM) compared to mock treatment. Methionine (Met), Homocysteine (Hcy), and oxidized glutathione (GSSG) were not tested (x) for the concentration of 10 μM. **C.** Boxplot showing the average biomass of individual plants treated with either mock solution (sterile water) or 100 nM GSH in both S-sufficient and S-deficient conditions. Number of biological replicates, n = 70 across three independent experiments, N = 3 (indicated by shape of data points). Statistical significance (denoted by letters on top of the boxes) was determined using ANOVA, controlling for experiments, followed by a post-hoc Tukey’s test (q < 0.05). **D.** Bar plot showing GSH concentration in both intra (top panel) and extracellular (bottom panel) fractions of *Pseudomonas sp.* X, *wild-type* and deletion knockouts of gshA (ΔgshA) strains. A standard curve using the peak area of quantifier ion (parent and product ions m/z = 308.1 and 179, respectively) and known GSH concentrations was used to calculate the absolute concentration of GSH in the extracellular and intracellular fractions (Table S16). Data are shown for mean ± standard error of mean (SEM), and the number of biological replicates is indicated on top of each bar. Statistical significance is calculated with t-test (p < 0.00001***). Representative extracted ion-chromatograms (EICs) from multiple reaction monitoring (MRM) data of GSH quantifier ion (308.1 > 179.0) in both intracellular (top panel) and extracellular fractions (bottom panel) of X (black) and ΔgshA (red). **E.** Representative image of 12-days old Arabidopsis shoots treated with cell-free extracts of X and ΔgshA strains in both S-sufficient and S-deficient conditions. Scale bar represents 2 cm length. **F.** Boxplot showing the relative increase in shoot area by cell-free extract from X (purple boxes and data points) and ΔgshA strains (turquoise boxes and data points) under both S-sufficient and S-deficient conditions. Number of biological replicates, n = 59 across three independent experiments (N = 3 indicated by shape of the data points). Statistical significance (denoted by letters on top of the boxes) was determined using ANOVA, controlling for experiments, followed by a post-hoc Tukey’s test (q < 0.05).

Given the co-functioning of S-metabolism between the plant and rhizosphere bacterial communities (Figure 2C), we hypothesized that the bacterial bioactive molecule might be S-containing (but not in the form of sulfate) and functions either through S provisioning or signaling. To test this hypothesis, we screened several known S-containing compounds, guided by knowledge of S-metabolic genes that were differentially abundant in the rhizosphere metagenomes of S over-accumulating plant genotypes identified earlier (Figure 2C). The screening of these chemicals was conducted at physiologically relevant concentrations ranging from micromolar to nanomolar levels, to represent S provisioning and signaling concentrations, respectively (Figure 6B). Among the tested chemicals, cysteine and glutathione showed the greatest ability to rescue plant growth under S-deficient condition, with glutathione (GSH) showing the highest effect size at 100 nM. This finding suggests that GSH may act as a potential bioactive chemical signal from rhizosphere bacteria (Figures 6B and 6C).

To validate the role of bacterial GSH as a potential signal for S-deficiency tolerance in plants, we selected a representative strain from SPAF18, *Pseudomonas* sp. X. This strain was selected for its ease of genetic manipulation and was among the top performing strains to impart S-deficiency tolerance in plants (Figures 4A, S5A, and S5B). A mutant derivative of this strain carrying deletion of the glutathione biosynthetic gene *gshA* (ΔgshA) accumulated negligible levels of GSH in both intracellular and extracellular fractions compared to the *wild-type* (WT) strain (Figure 6D and Table S16). Notably, GSH was negligible in the extracellular fraction of ΔgshA, whereas it was detected at nanomolar concentrations (53.36 ± 3.07 nM) in the exudates of the WT strain. This exudate from the WT strain was able to impart S-deficiency tolerance by increasing the shoot area under S-deficient condition (Figures 6D, 6E, and 6F), supporting our finding that GSH is bioactive at nanomolar (signaling) concentration (Figure 6C). In contrast, ΔgshA failed to enhance shoot area under S-deficiency, confirming that GSH is essential for its beneficial effect (Figures 6E and 6F). Together, these results confirm the role of bacterial GSH as a novel ‘cell-free’ and ‘water-soluble’ signal for S-deficiency tolerance in plants.

Since GSH is required by both plants and the rhizosphere microbiome, we further investigated if stronger competition among bacterial strains releases more GSH, according to our proposed ‘trans-kingdom fitness’ model. We obtained a moderate and marginally significant correlation (r = −0.44; p = 0.06; Figure S7D and Table S17) between the strength of bacterial competition and the amount of GSH released in a bacterial pair. This supports our tradeoff model that stronger bacterial competition (calculated as sum of interaction scores in a bacterial pair) releases more GSH in their extracellular space, thereby making it available to the plants.

## Discussion

We report here an interkingdom phenomenon in which the rhizosphere microbiome alleviates S-deficiency responses in plants via an extracellular and water-soluble signal, GSH. Previous reports have identified the role of microbiome in provisioning nitrogen and iron in their bioavailable forms in response to plant chemical signals.^32,33^ In the case of S, volatile DMDS from isolated *Bacillus* sp. B55 has been shown to alleviate S-deficiency responses in tobacco plants.^27^ However, alleviation of S-deficiency through services of the rhizosphere microbiome has not been reported previously. Given that S is utilized as a plant nutrient in its diverse forms due to its flexible redox states, we had investigated the relationships between different plant S nutrition and the rhizosphere microbiome. A combination of chemical and genetic approaches revealed that GSH is a central signaling molecule in this communication between the rhizosphere microbiome and the host plants. We have shown that GSH is bioactive in the nanomolar range, as demonstrated in our supplementation experiment and its quantification in bacterial cell-free extract (Figures 6C and 6D). Hence, at nanomolar levels, we can discount provisioning as its likely mode of action. GSH, which is essential across all domains of life including plants, bacteria, and fungi, is well-recognized for its role in mitigating oxidative stress, bacterial pathogenesis, biofilm formation, and cell cycle progression, among many others.^34,35,36^ Our findings reveal a previously unrecognized role of GSH within the exo-metabolome of the rhizosphere microbiome that improves plant growth under S-deficiency. Plants generate reactive oxygen species (ROS) in response to S-deficiency, which leads to oxidative stress.^37,38,39^ Counteracting this stress is essential for bacterial colonization within the root environment, ^40,41^ which has been shown to require GSH.^42^ Our findings suggest that GSH may have been co-opted from the rhizospheric microbes to counteract ROS toxicity and signal the plants to develop S-deficiency tolerance. This is supported by our observation that cell-free extracts from an isolated strain with loss of GSH biosynthetic gene is unable to rescue plant growth under S-deficient condition compared to the extracts from its *wild-type* strain that can produce GSH. An independent study has further established the role of exogenous GSH in mitigating ROS toxicity in plants.^43^

The overaccumulation of sulfate in selected genotypes (*sdi2;1* and *msa1-3*) resulted in significant alterations in composition and functions (particularly S-metabolism) of the rhizosphere microbiome. This prompted us to develop a SynCom that represented the S-metabolic potential of the rhizosphere microbiome members (Figures S3C, S3D, and S3E). This potential was realized by the 18-membered SynCom which imparted tolerance towards S-deficiency via a S-containing signaling molecule, GSH (Figure 3). This function-based approach to assemble SynCom members marks a significant departure from previous methods, which often rely on random assembly or phylogenetic representation from a culture-based collection.^44,45^

Microbial interactions lead to emergent properties that significantly influence host fitness.^46,47,48^ In our context of host fitness towards S-deficiency tolerance, both computational predictions based on genome-scale metabolic models (Figures 5B and S7B) and experimental validations (Figure S7C and Table S14) showed diverse microbial interactions-amensal, competitive, and neutral between bacterial pairs. Among these, competitive interactions were the vast majority, which underscores their prevalence in microbial communities, which is also supported by other studies.^49,50,51^ But how do these interactions influence plant fitness? Interestingly, stronger competitive interactions between bacterial pairs led to greater fitness benefits to the host plants under S-deficient conditions. We propose that this phenomenon is an example of ‘trans-kingdom fitness tradeoff’, wherein the host benefits at the expense of fitness of host-associated bacterial members. We present one such possible mechanism wherein stronger bacterial competition (calculated as sum of interaction scores in a bacterial pair) releases more GSH in their extracellular space, making it available to the host plant. Tradeoffs of this sort are probably more widespread across diverse host-microbiome (holobionts) systems and may involve multiple mechanism(s), other than through GSH, as presented here.

To understand whether the effects of bacterial interactions on plant fitness can be explained through the above pairwise combinations or there could be community-level interactions giving rise to additional emergent properties, machine-learning based predictive models could be highly useful. As a first step in this direction, we assessed the suitability of 15 different machine learning models for their ability to predict the plant benefits starting from the simplified case of pairwise microbial interactions. Among the models tested for all possible 153 pairwise interactions, tree-based models performed significantly better than the others (Table S9). Clustering the bacterial members into modules (to reduce dimensions) significantly improved the model’s predictive performance in our dataset which is consistent with previous reports on the use of this approach to improve the predictive performance of such models.^52^ There is precedence in some host-associated communities where pairwise interactions, as tested here, can explain the behaviors of complex host-associated communities.^48,53^ However, our current study has a limitation that it addresses only pairwise interactions and not higher-order ones within the microbial community. It will, therefore, be interesting to extend the current tree-based machine learning models to predict higher-order interactions within the rhizosphere microbial communities.

In conclusion, this study establishes that the well-known antioxidant GSH is an extracellular chemical produced by the rhizosphere microbiome that is repurposed as a cross-kingdom signal to enhance the fitness of host plants under S-deficiency. This benefit is further amplified by a cross-kingdom fitness tradeoff that results from stronger competitive interactions releasing more GSH from the rhizosphere microbiome (Figure 7). This study advances our understanding of bacterial interactions and the chemical signals that mediate nutritional stress adaptation in plants, unveiling a unique phenomenon in the rhizosphere. Given the alarming increase in agricultural S applications, these findings hold significant implications for sustainable crop production in S-deficient agricultural soils worldwide.^16,17^ Molecular insights onto how bacterial competition leads to more GSH release and how the host plant perceives exogenous GSH, and its subsequent signal transduction mechanisms will deepen our understanding of this unique phenomenon. Furthermore, exploring the emergent functions arising from metabolic interactions in complex rhizosphere microbiomes will shed light on their ecological implications and inform strategies to deploy microbial inoculants for sustainable crop production.

**Figure 7.**
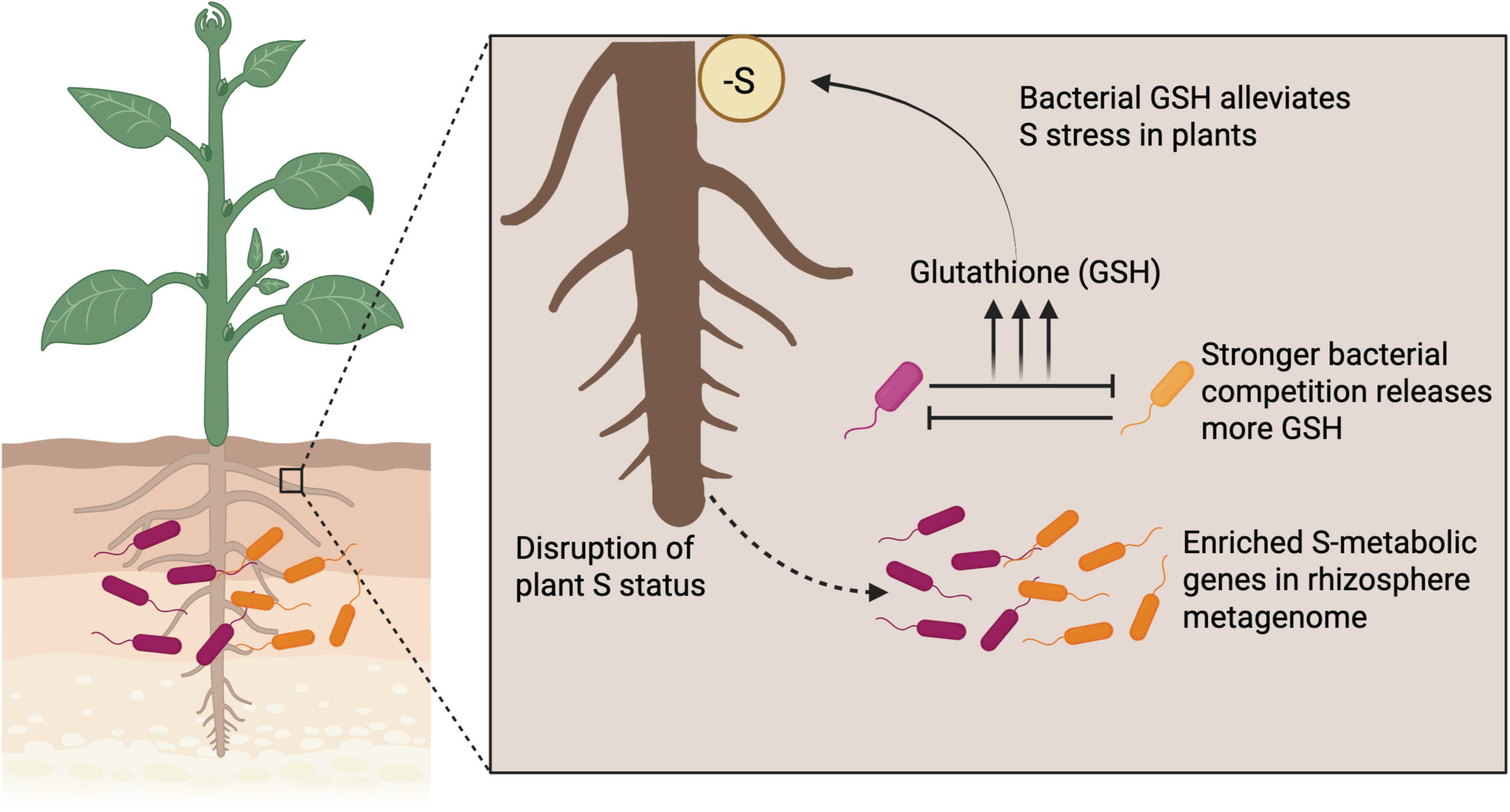
Conceptual diagram showing GSH mediated transkingdom fitness tradeoff to improve S deficiency tolerance in plants. Disruption in plant S content leads to enrichment of S-metabolic genes in the rhizosphere metagenomes. Stronger bacterial competition lead to enhanced GSH release which in turn signals the plant to improve S-stress tolerance.

### Limitations of the study

The role of GSH as an extracellular chemical signal was demonstrated using a single strain, *Pseudomonas sp.* X and all bacterial strains in our collection could release GSH in their extracellular space. However, to make broader claims, it will be important to evaluate GSH biosynthetic mutants in additional rhizosphere bacterial strains. Additionally, since we tested only a limited number of bacterial pairs for their effects on plant growth promotion under S deficiency, the predictive accuracy of our machine-learning model was moderate. Expanding the training dataset would likely enhance the predictive performance.

## Materials and Methods

### Sources of soil and soil preparation

We conducted our pot-based screening experiment using BVB Black Peat Moss (Holland) soil, which we procured from *Bioflora Ltd.* in Singapore. Upon receiving the soil, we stored it in clean plastic bags at 4°C until use. Before the experiment, we dried the soil in clean plastic trays for five days and sifted it using a plastic sieve. The soil was then thoroughly mixed with autoclaved sand in a 2:1 (v/v) ratio and sieved again to remove debris. Drying the soil was essential to ensure uniform moisture content, and pulverization was performed to avoid heterogeneity. The pots were sterilized with 70% ethanol to prevent unwanted bacterial contamination. We used disinfected pots (11×11×10 cm), filled them with the prepared soil mixture, and used them to grow Arabidopsis.

### Plant materials and growth condition

All insertional mutant lines (Col-0 background) were obtained from the Arabidopsis Biological Research Centre at Ohio State University (ABRC: http://www.arabidopsis.org/abrc/; Table S1). The seeds were grown in Black Peat Moss soil and screened for homozygous T-DNA insertion lines according to instructions from ABRC. The selected lines were then multiplied for seed collection, dried, and stored at ambient temperature for subsequent experiments.

Seeds were surface-sterilized with 50% bleach for 5 minutes and rinsed with sterile distilled water at least five times to eliminate any seed-borne microbes. They were stratified at 4°C in the dark for 3 days and then germinated in sterile pots filled with soil prepared as described before. As controls, we used pots without plants as ‘bulk soil’ and pots with wild-type Col-0 plants. All pots were randomized using a true random generator (random.org), and trays were reshuffled every week in the growth chamber without regard to pot labels. All pots were watered twice a week by filling the bottom tray and spraying the top with sterile distilled water to avoid chlorine and other tap water additives. Plants were grown in a growth chamber with a 16-hour light / 8-hour dark regime at 21°C for 5 weeks. Due to poor germination rates of *sultr2;1* seed, we included them only in the second replication of our pot-based study.^54^ A small-scale experiment with the genotypes Col-0, *sdi2;1*, and *sultr1;1* is also included.

### Anion analysis

Shoots of five-week-old plants were collected using sterile forceps, snap-frozen in liquid nitrogen, and stored at −80°C until further use. The shoots were then freeze-dried, pulverized using stainless steel beads, and weighed (10-12 mg) for anion analysis. The dried powder was reconstituted in 1 mL of sterile distilled water, sonicated three times at maximum speed, and centrifuged to remove debris. The supernatant was filtered through a 0.2 µm filter, and the volume was adjusted to 10 mL before being submitted for anion analysis (nitrate, phosphate, and sulfate) at the Elementary Analysis Laboratory (EAL), Department of Chemistry, National University of Singapore. The concentration of each sample was provided in parts per million (ppm) and then converted to µmol per mg of dry matter using the following formula:

Anion content (µmol per mg of dry matter) = (Concentration in ppm x 1000) / (dry weight in mg x M);

M is the Molecular weight of the anion (nitrate: 62.0049, phosphate: 94.9714, or sulfate: 96.07).

To test for differences among the genotypes, we first checked data distribution and homogeneity of variance using the Shapiro test and Levene’s test, respectively.^55,56^ Based on these results, we chose an appropriate statistical test (ANOVA or Kruskal-Wallis) to determine statistical significance (Benjamini-Hochberg corrected p-value < 0.1).

### Calculation of Cohen’s effect size

Cohen’s effect size (ES), which is a measure of difference between the *wild-type* Col-0 plants and the genotypes for anion content (sulfate, phosphate, and nitrate) and shoot biomass. The following formula was used to calculate Cohen’s ES:

Cohen’s ES = (M_2_ - M_1_) ⁄ SD _pooled_
M_1_ = Sample mean for mutant
M_2_ = Sample mean for *wild-type* Col-0
SD _pooled_ = √ ((SD_1_^2^ + SD_2_^2^) ⁄ 2), where SD_1_ and SD_2_ are the standard deviations for sample group 1 and 2. Here, sample group 1 refers to the mutant group samples and sample group 2 refers to the *wild-type* Col-0 samples.

### Microbiome profiling

We analyzed the microbial community composition of the root, shoot, and rhizosphere from samples of five genotypes (Figure 1B) using 16S amplicon sequencing. On the day of sampling, each pot containing a five-week-old single plant was inverted to remove the soil manually until only a fine layer of soil remained attached to the roots. The shoots and roots were separated using a sterile blade and placed in sterile water and Phosphate Buffer Saline (PBS) in 50 mL tubes, respectively.

The shoots in sterile water and the roots (with the thin layer of attached soil) in PBS were sonicated for 5 minutes at maximum frequency to remove weakly adhering epiphytes from the shoots and to obtain rhizosphere soil from the roots. The washed roots and shoots were then dried, placed in sterile 2 mL tubes containing sterile steel beads, snap-frozen in liquid nitrogen, and stored at −80°C until further use. Rhizosphere soil samples in PBS were centrifuged at 8,000 x g for 5 minutes to remove the supernatant, snap-frozen, and stored in the same manner as the root and shoot samples.

DNA extraction for root and shoot samples was performed according to the manufacturer’s protocol using the Qiagen DNeasy Plant Mini Kit (Catalog No. 69104). For rhizosphere samples, we followed the protocol provided in the Zymo Quick-DNA Fecal/Soil Microbe Miniprep Kit (Catalog No. D6010). The extracted DNA was quantified using the Qubit DNA BR assay kit and stored at −20°C until further use.

The concentration of the amplicons was quantified and diluted to 10-15 ng / µL of DNA when applicable. We also included a no-template negative control and a known synthetic community DNA (positive control) along with the actual samples to remove spurious sequences and estimate sequencing accuracy. Primers for 16S amplicon profiling can be found in Table S2. The 16S amplicons were then submitted for sequencing on an Illumina MiSeq V3 platform (300 bp paired-end) at the Singapore Centre for Life Sciences Engineering (SCELSE).

### Processing of raw amplicon sequences

Raw and demultiplexed sequences were quality-trimmed using Cutadapt^56^ to remove primers, adapters, and low-quality bases (bases with a Q-score <30 were trimmed). The high-quality reads were then processed with the DADA2^57^ pipeline to obtain an Amplicon Sequence Variant (ASV) table, and the ASVs were assigned taxonomy based on the SILVA database (v138.1).^58^ ASVs assigned as chloroplast, mitochondria, or unassigned were removed prior to statistical analysis.

### Amplicon data analysis

The ASV table, taxonomy table, and metadata information were processed in R to infer alpha and beta diversity separately for shoot, root, and rhizosphere samples. Before computing diversity indices, ASVs with fewer than 10 total reads across samples and samples with insufficient diversity (less than 10,000 reads for root samples and less than 3 shannon diversity for shoot samples) were excluded. Next, we rarefied the samples from root, shoot, and rhizosphere to their minimum read depth. We also included a synthetic community with a known relative abundance of species, which was sequenced to check the accuracy of the amplicon sequence data processing (Figure S1E).

Alpha diversity was calculated using the Shannon diversity index from the vegan package in R.^59^ To test for differences in alpha diversity among genotypes, we used the similar statistical approach as described for anion content and shoot biomass data. To compare beta diversity across genotypes, we performed constrained ordination (constrained by Experiment and batch of PCR) based on Bray-Curtis dissimilarity among samples, using the capscale function from the vegan package in R. Statistical significance was assessed using PERMANOVA. To calculate beta diversity differences of the genotypes against Col-0, we performed pairwise PERMANOVA. The R script for detailed alpha and beta diversity analyses can be found at https://github.com/arijitnus/Nutrition_axis/tree/main/16S_microbiome.

Co-occurrence networks were prepared based on Spearman correlation using the MENA workflow.^60^ The ‘meconetcomp’ package in R was used for this with default parameters.^61^

### Metagenome sequencing

We performed metagenome sequencing on rhizosphere samples from Col-0, *msa1-3*, and *sdi2;1* genotype, with four samples for each genotype (n = 4). These were the same samples previously used for 16S amplicon sequencing, except for one randomly excluded sample from the Col-0 genotype. DNA was extracted from the rhizosphere soil samples using the same protocol as the 16S amplicon analysis. The total DNA was then submitted for sequencing on an Illumina HiSeq platform (250 bp paired end) at the Singapore Centre for Environmental Life Sciences Engineering (SCELSE). To obtain a higher coverage of metagenome reads for assembly, we used a subset of these samples (n = 4 each from Col-0 and *msa1-3* from Experiment 1) for sequencing on HiSeqX platform (150 bp paired-end).

### Processing of raw metagenome sequences and contigs assembly

Raw de-multiplexed reads were processed to remove adapters and trim low-quality regions using Cutadapt v2.3 and BBDuk v38.56, respectively.^56^ The quality-trimmed reads were then assembled both individually and through co-assembly using MEGAHIT v1.2.8.^62^ Contigs shorter than 1 kbp were discarded. Read containment was estimated by mapping the quality-trimmed reads to the assembled contigs using Bowtie2 v2.3.5.^63^ The parameters used for each of these tools are detailed in our previous publication.^9^

### Prediction of functional potential in metagenomes

We predicted the functions of the assembled contigs using Cluster of Orthologous Genes (COGs) and KEGG, with Prokka and Kofamscan respectively, both with default parameters.^64,65^ The count of predicted genes for each sample was compiled into a table and used for subsequent statistical analyses.

### Metagenome data analysis

To understand the overall differences in the functions of rhizosphere metagenomes, we performed constrained ordination using the ‘cao’ distance measures among the samples. This was done with the ‘capscalè function from the ‘vegan’ package in R, and statistical significance was assessed using PERMANOVA.

To determine the differential representation of genes in rhizosphere metagenomes of the different genotypes, we combined functions from both COG and KEGG analyses and performed differential analyses. We used the ‘metagenomeSeq’ package in R to identify differentially represented genes in the metagenomes of *sdi2;1* and *msa1-3* genotypes compared to Col-0 samples, following the method described previously.^66^ Briefly, the count tables were first normalized using the ‘cumNorm’ function, which performs cumulative sum scaling (CSS) normalization. Next, we fitted a generalized linear model using the ‘fitZig’ function from metagenomeSeq and conducted moderated t-tests with the fitted coefficients to calculate p-values. Statistically significant COGs (FDR corrected p-value or q < 0.1) were considered to identify GO Biological processes that are differentially represented in these genotypes. COG IDs were mapped to COG categories, and over-representation of COG categories were calculated based on Fisher’s Exact test (adj-P < 0.1). The proportion of genes differentially represented and corresponding q values were reported (Figure 2B and Table S5). To visualize differentially represented genes, we have only considered the sulfur metabolism related genes from both KEGG and COG. We calculated their scaled average counts (per million mapped reads) across biological replicates within each genotype (Figure 2C).

### Genome binning, decontamination, dereplication, and quality assessment

Contigs longer than 1 kbp were assembled into bins using MetaBAT2 and MaxBin.^67,68^ The bins from these tools were then aggregated and de-replicated using DAS Tool.^69^ Refinement of the bins was done by removing contigs with divergent genomic properties or taxonomy using RefineM and MAGPurify, respectively.^70^ Assembly statistics for the MAGs, including completeness, redundancy, size, number of contigs, contig N50, length of the longest contig, average GC content, and the number of predicted genes, were computed using the lineage workflow from CheckM v1.0.18.^71^ Taxonomy assignment of the MAGs was performed using GTDB-Tk.^72^ MAGs were classified as low, medium, or high-quality based on their completeness and contamination (High quality: completeness >90% and contamination <10%; medium-quality: completeness 50-90% and contamination <10%; low-quality: completeness <50% or contamination >10%). Summary statistics for all the MAGs are provided in Figure S3A. The parameters used for each tool are detailed in our previous publication.^73^

### Bacterial culture collection and development of synthetic rhizosphere bacterial community

To assemble a synthetic rhizosphere bacterial community (SynCom), we utilized an in-house collection of cultured bacteria from the rhizosphere of field grown crops-ChoySum (*Brassica rapa var. parachinensis*), Bayam (*Amaranthus tricolor*), and Kai lan (*Brassica oleracea* var. *alboglabra*), from *Kok Fah Technology Farm Pte Ltd*, Singapore (1.3938° N, 103.7280° E). From the initial 55 genome-sequenced bacterial isolates, we first identified pure and non-redundant strains, selected strains with 16SrRNA identity of less than 97%, and checked their ability to grow till saturation in ¼ TSB (Tryptic Soy Broth). The resulting 18 genomes possessed at least one unique S-metabolic gene; therefore, we didn’t further reduce the community to represent the maximum number of S-metabolic genes. This 18-member synthetic community which we named as SPAF18 (<u>S</u>ulfur <u>P</u>roteobacteria <u>A</u>ctinobacteria <u>F</u>irmicutes 18) (Figure S4A).

### Functional potential of SPAF18 and MAGs

We predicted sulfur metabolic genes for both MAGs and SynCom members from KEGG to compare their functions. To compare the representation of sulfur metabolic genes between MAGs and SynCom members, we used unconstrained ordination based on ‘Bray-Curtis’ distance measures (Figure S4B). We also compared the representation of sulfur metabolic genes of SPAF18 with a global collection of rhizosphere microbiomes, as published previously (Figure S4C).

### SynCom experiments with Arabidopsis under sulfur stress in plate-based gnotobiotic systems

#### Plant growth condition

Surface sterilization and vernalization of seeds were performed as described previously. Seeds were germinated on MS-agar (with 1% sucrose) for 6 days and transferred to sulfur sufficient/deficient media containing plates. Sulfur-deficient (S-deficient) media was prepared by replacing MgSO_4_ with MgNO_3_. For S-sufficient media, we added 1500 µM sodium sulfate to the S-deficient media. A detailed composition of the S-sufficient and S-deficient media is provided in Table S11.

#### Bacterial culture and plant inoculation

One day before the experiment, overnight grown cultures of SPAF18 with a starting OD of 0.02 were inoculated in 1⁄4 TSB. We prepared two copies of each of the SPAF18 members to represent both, metabolically active (4h) and stationary phase (24h) of cells. Cells were washed twice with 10 mM MgCl_2_ and resuspended to a final OD of 0.1 in 10 mM MgCl_2_. All members were mixed in equal volumes to prepare the final SPAF18 community, and 50 µL of this was spread in each plate (sulfur sufficient or deficient) before placing 4days old seedlings (5-6 seedlings per plate). Plates were kept in a 16-hour light/8-hour dark regime at 21°C for seven days before sampling. Mono-association and co-inoculation experiments were performed similarly.

### Plant phenotypes

We estimated five plant parameters-primary root length, root network, number of lateral roots, shoot area, and biomass. These parameters were estimated using ImageJ with a custom in-house macro, as mentioned in our previous publication.^5^ The Cohen’s effect size of the active SynCom was calculated for each phenotype, relative to their heat killed controls, as described above for Figure 1D. Detailed datasets are provided in Table S8.

### SynCom experiments with ChoySum plants

We tested the applicability of SPAF18 in a Brassicaceae crop ChoySum, locally grown in Singapore. ChoySum seeds were sterilized and vernalized using a similar protocol as described above for Arabidopsis. Following germination after 5 days, we transferred individual plants in vermiculite filled plastic pots with either S-sufficient or S-deficient media (around 70 mL per pot) with either active or heat-killed SPAF18. Plants were grown for an additional 10 days before sampling and phenotyping.

### Mono-association and co-inoculation experiments

Mono-association and co-inoculation experiments were performed using the same protocol for SPAF18 inoculation.

### Genome scale metabolic models and pairwise interactions

We constructed genome-scale metabolic models for all members of SPAF18 using gapseq.^74^ This tool predicts the Reactions, pathways, and transporters from genome sequence FASTA files. Using these draft models, we predicted their completed metabolic model by ‘Gapfilling’ algorithm in Artificial root exudates + MS + Vitamins media, as this closely resembled our experimental setups (Table S11). Growth rates were simulated with exchange rates adopted from a previous study and can be found in Table S10.^49^ Quality metrics of the models were checked using MEMOTE and are available in Table S10.^75^ Two of the strains (*Priestia* sp. 7F21 and *Paenibacillus* sp. Y) did not pass the QC and therefore were removed from further analyses. We predicted their pairwise interactions based on their growth rates in monoculture and in co-culture with the interacting strain. If the growth rate of the focal strain was higher or lower by 10% in co-culture with the interacting strain than its growth rate in monoculture, we considered that interaction to be positive and negative, respectively.

### Validation of pairwise bacterial interactions

We validated pairwise bacterial interactions using CFU counting. *Pseudomonas* sp. X is autofluorescent in King’s B agar, and therefore, allows easier distinction with other strains. Selected strain pairs were grown in Artificial root exudates + MS media (Table S13) with equal starting OD of 0.01 for both monoculture and coculture. Following growth for 24 hours, we prepared dilutions ranging from 10^-4^ to 10^-8^. 30 µL of these dilutions were mixed with 70 µL of sterile 10 mM MgCl_2_ and plated on King’s B agar plates and incubated at 30°C for 24h. CFUs in those plates were counted using ImageJ and interaction scores were calculated using the following formula:

Interaction score of strain A in coculture with strain B = Log_2_ (CFUs of strain A in monoculture / CFUs of strain A in coculture)

### Machine learning models

We used both mono association and pairwise combination data for preparing the ML models. Average increase in biomass relative heat killed control (n = 6) were used to train the ML models. First, we clustered the individual bacterial strains based on their ability to influence plant phenotypes into five clusters (Figure S7A). This approach of clustering the individual strains helps avoid the ‘curse of multidimensionality’ ^76^. Individual strains within each cluster were assigned relative importance based on their performance within each cluster (Strains with the highest relative biomass promotion were assigned the highest score of 1). Subsequently, pairwise interaction data were considered for training the ML models. Each pair was assigned weights based on their origin of clusters and indexed. If two bacteria were selected from the same cluster, their weights were summed up. We fitted 15 machine learning models and tuned their hyperparameters to find the best performing models. Random forest regression model performed best among the models tested. Performance of the models were tested based on R^2^, root mean square error (RMSE), and mean absolute error (MAE) of training and test data (67:33 split). Many of the models (non-tree based) overfitted, and we observed relatively better performance of the tree-based models (Table S8). For the best performing model-random forest regression, we predicted the relative increase in biomass for all pairwise combinations (^18^C_2_ = 153).

### Cell-free extract experiment

SPAF18 members were grown overnight in ¼ TSB media, centrifuged at 8,000 x g for 5 mins and filtered through 0.22 μm membrane to remove bacterial cells. This cell-free extract was combined in equal volume from SPAF18 members and spread onto plates (70 µL) before planting 4d old seedlings.

### Chemical supplementation experiments

We selected six S-containing chemicals (from Sigma Aldrich) for this screening at different concentrations (ranging from nM to mM). Except for Cysteine, all other stock solutions of these chemicals were prepared in MilliQ water, filter-sterilized and spread onto plates (70 µL) before planting 4-days old seedlings. Cysteine stock solution was prepared in 1 M HCl and then diluted with MilliQ water to prepare their respective concentrations and 70 µL was spread on agar plates before planting 4-days old seedlings.

### Bacterial deletion mutant generation

We generated marker-free knockout for glutathione biosynthetic gene in *Pseudomonas sp*. X. GSH biosynthesis is a two-step catalytic process, catalyzed by glutamate-cysteine ligase (*gshA*) and glutathione synthetase (*gshB*). The former catalyzes the bonding between glutamate and cysteine to form γ-L-glutamylcysteine, which is then converted into GSH by the latter.

Knockout was generated for *gshA* through homologous recombination using the cloning vector pk18 containing genes for resistance to Gentamicin and susceptibility to sucrose (*sacB* gene). ∼500 bp upstream and ∼1 kb downstream of the target genes were cloned into this plasmid through Gibson assembly. The resulting plasmid construct was transformed into *E. coli* S17-1 λpir strain and used for conjugation with *Pseudomonas sp*. X. The plasmid is integrated into receiving strain through homologous recombination and deletion mutants were generated by second counter-selection mediated homologous recombination. The primers used for generating deletion knockout strain can be found in Table S2.

#### Cloning of pk18-derived deletion construct for target gene

∼500 bp upstream and ∼1 kb downstream region of the target gene was amplified with primers flanking regions of the pk18 plasmid. These fragments were ligated with pk18 using Gibson assembly. The ligated product was confirmed with sanger sequencing.

#### Biparental conjugation

Sequence confirmed constructs were transformed into *E.coli* S17-1 λpir strain (donor) for biparental conjugation with *Pseudomonas sp*. X (receiver). Briefly, Donor and receiver strains were grown in LB containing 30 µg/mL Gentamicin sulfate and LB broth, respectively at 37°C for 6-7 hrs. We measured the optical density (OD) of both the strains and took an equal number of cells in 500 µL. The cells were centrifuged at 2000 x g for 8 mins and washed with sterile 10 mM MgSO_4_. The cell pellets were then resuspended in 50 µL LB media and mixed. The 100 µL mix of donor and receiver cells were spotted on LB agar plates, dried and incubated at 30°C for 20-24 hrs to allow conjugation events.

#### Screening for single recombinants

The conjugation mix was resuspended in 4 mL ABTC media with sterile loops. 100 µL of the mix was plated in ABTC agar plates containing 30 µg/mL Gentamicin sulfate. This mix was incubated at 37 °C for 48 hrs. Colonies were screened for single recombinants with plasmid specific and gene specific primers.

#### Counter selection with sucrose for double recombinants and mutant confirmation

Confirmed single recombinants were grown in ABTC broth for 2 hours until they became slightly turbid, facilitating double crossovers. Following this, 100 µL was plated on ABTC agar plates containing 5% sucrose to allow for selection of double crossovers with deletion strains. Single colonies growing on ABTC agar plates (with 5% sucrose) were spotted onto ABTC agar plates and ABTC agar plates containing 30 µg/mL Gentamicin sulfate. Sucrose resistant and Gentamicin sensitive colonies were further confirmed for deletion with PCR using primers flanking the PCR products used to generate deletion. The PCR products were gel purified and sent for sanger sequencing for confirmation of deletion.

### Bacterial metabolites extraction and quantification of GSH

*Pseudomonas* sp. X and the deletion strains were grown in 5 mL M9 media (Supplemented with glucose as the carbon source) overnight. Saturated cultures were centrifuged at 8,000 rpm for 5 mins, washed with sterile PBS and 1 mL of 0.5 O.D. of individual cultures were taken for intracellular metabolites extraction. Briefly, cell pellets were resuspended in 1 mL of 50% MeOH and sonicated for 20 mins on ice. The solution was passed through a 0.22 µm membrane to remove any debris. The solution was injected in QQQ for targeted identification of GSH. To prepare the exo-metabolites fraction, we filtered the 5-10 mL of spent media (adjusted to 1 O.D of the overnight culture) through a 0.22 µm membrane, freeze-dried, and reconstituted in 1 mL 50% MeOH before performing targeted metabolomics.

Samples were run on Agilent Triple-quad (QQQ) with the following gradient: 20% ACN (Acetonitrile) for 2 mins, 80% ACN for 5 mins followed by washing. We used ACQUITY^TM^ PREMIER (Waters) HSS T3 column (1.8 µm x 2.1 mm x 100 mm) for this analysis. A standard curve based on known concentrations of GSH (from 10 ppb to 500 ppb) was used for calculating concentration of GSH in intracellular and extracellular fractions. Briefly, the transition ion (parent ion m/z 308.1 > product ion m/z 176.0) was used as the quantifier ion. Peak area for this transition pair was used from the known GSH concentrations (10 ppb to 500 ppb) to prepare a standard curve. Sample concentration was then measured from the linear regression equation of the standard curve and converted to nanomolar (nM) to report their absolute concentration.

## Author contributions

Conceptualization, A.M., and S.S.; Methodology, A.M., M.M., A.V, R.B., and S.S.; Investigation, A.M., M.M., A.V., and O.Q.E.; Writing– Original Draft, A.M.; Writing– Review & Editing, A.M., M.M., and S.S.; Funding Acquisition, S.S.; Resources, S.S.; Supervision, S.S.

## Acknowledgements

This research was funded by the Singapore Centre for Environmental Life Sciences Engineering (SCELSE). We thank SCELSE for the Ph.D. research scholarships to A.M. We thank the National University of Singapore for providing us the lab facilities necessary for conducting our research. We thank Dr Sujatha Subramoni and Dr Goh Yu Fen for passing the *E.coli* S17-1 λpir strain and sharing the protocol for deletion strain generation. We also want to thank Mr. Sevagun Mayalagu for assistance during sampling and Mr Boon Hao Tan for help in CFU based experiments. The authors also acknowledge the use of the recently developed ‘ChatGPT’ for R codes, grammatical corrections, and improved phrasing in this manuscript.

## Resource availability

### Lead contact

Further information and requests for resources and reagents should be directed to and will be fulfilled by Sanjay Swarup.

### Materials availability

All plant mutants used in this publication are available at ABRC (Arabidopsis Biological Resource Center) with the accessions provided in Table S1. All bacterial strains are available upon request to the lead contact.

### Data and code availability

The raw sequencing data have been deposited in the Sequence Read Archive (SRA BioProject: PRJNA1114376 for rhizosphere and root samples; SRA BioProject: PRJNA1114383 for shoot samples). Raw metagenome sequencing reads are available with accession SRA BioProject: PRJNA1177226. Raw metabolomics data for targeted quantification of GSH are available upon request to the lead contact. All codes to reproduce the analyses reported in this article has been deposited at GitHub and is publicly available at https://github.com/arijitnus/Nutrition_axis.

## Supplemental information

**Document S1.** Figures S1-S7

**Tables S1-S17.** Excel file describing the experimental and extended datasets related to Figures 1-7.

