## Supplementary Figures for "A cell-free bacterial signal orchestrates trans-kingdom fitness tradeoff to enhance sulfur deficiency tolerance in plants"

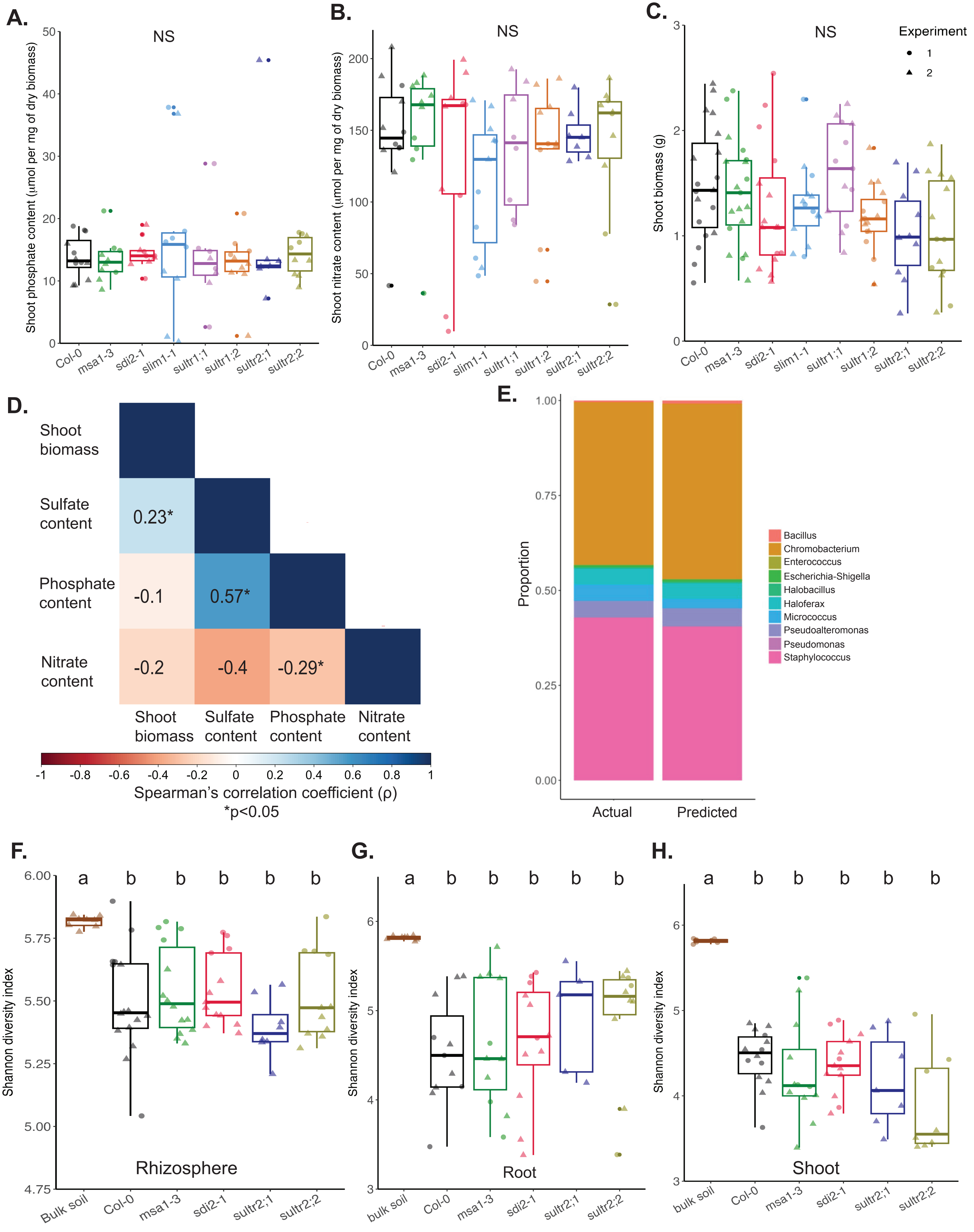
A.
NS
Shoot phosphate content (µmol per mg of dry biomass)
Col-0
msa1-3
sdi2-1
slim1-1
sultr1;1
sultr1;2
sultr2;1
sultr2;2
B.
NS
Shoot nitrate content (µmol per mg of dry biomass)
C.
NS
Experiment
1
2
Shoot biomass (g)
D.
Shoot biomass
Sulfate content
Phosphate content
Nitrate content
0.23*
-0.1
0.57*
-0.2
-0.4
-0.29*
Spearman's correlation coefficient (ρ)
*p<0.05
E.
Proportion
Actual
Predicted
Bacillus
Chromobacterium
Enterococcus
Escherichia-Shigella
Halobacillus
Haloferax
Micrococcus
Pseudoalteromonas
Pseudomonas
Staphylococcus
F.
Shannon diversity index
Rhizosphere
Bulk soil
G.
Root
bulk soil
H.
Shoot
a
b

**Figure S1. Anion content, shoot biomass, and microbiome composition of the genotypes considered in this study**

**A.** Boxplot showing phosphate content for all genotypes (Col-0: n = 12; *msa1-3*: n = 10; *sdi2;1*: n = 10; *slim1-1*: n = 10; *sultr1;1*: n = 8; *sultr1;2*: n = 10; *sultr2;1*: n = 7; *sultr2;2*: n = 10). **B.** Boxplot showing nitrate content for all genotypes (Col-0: n = 12; *msa1-3*: n = 10; *sdi2;1*: n = 10; *slim1-1*: n = 11; *sultr1;1*: n = 8; *sultr1;2*: n = 10; *sultr2;1*: n=7; *sultr2;2*: n=10). **C.** Boxplot showing shoot biomass for all genotypes (Col-0: n=19; *msa1-3*: n=19; *sdi2;1*: n = 16; *slim1-1*: n = 14; *sultr1;1*: n = 13; *sultr1;2*: n = 16; *sultr2;1*: n = 10; *sultr2;2*: n = 14). Data points are shown from two independent experiments (indicated by shape of points) and colored according to the genotypes. **D.** Correlation matrix showing the pairwise correlation coefficient (Spearman's $\rho$) for anion contents and shoot biomass. Statistically significant correlations are indicated with an asterisk ($p < 0.05$*). **E**. Stacked bar plot showing the relative abundance of a known synthetic community and the same community predicted through 16S amplicon sequencing. **F, G, and H.** Shannon diversity index of rhizosphere, root, and shoot microbiome of selected genotypes. Data points are shown from two independent experiments (indicated by shape of points) and colored according to genotypes. Number of biological replicates: bulk soil (n = 7), Col-0 (n = 16,11, and 14 in rhizosphere, root, and shoot, respectively), *msa1-3* (n = 13,11, and 12 in rhizosphere, root, and shoot, respectively), *sdi2;1* (n = 13,12, and 13 in rhizosphere, root, and shoot, respectively), *sultr2;1* (n = 8,5, and 7 in rhizosphere, root, and shoot, respectively), and *sultr2;2* (n = 11,10, and 8 in rhizosphere, root, and shoot, respectively), distributed across two independent experiments (indicated by shape of the points).

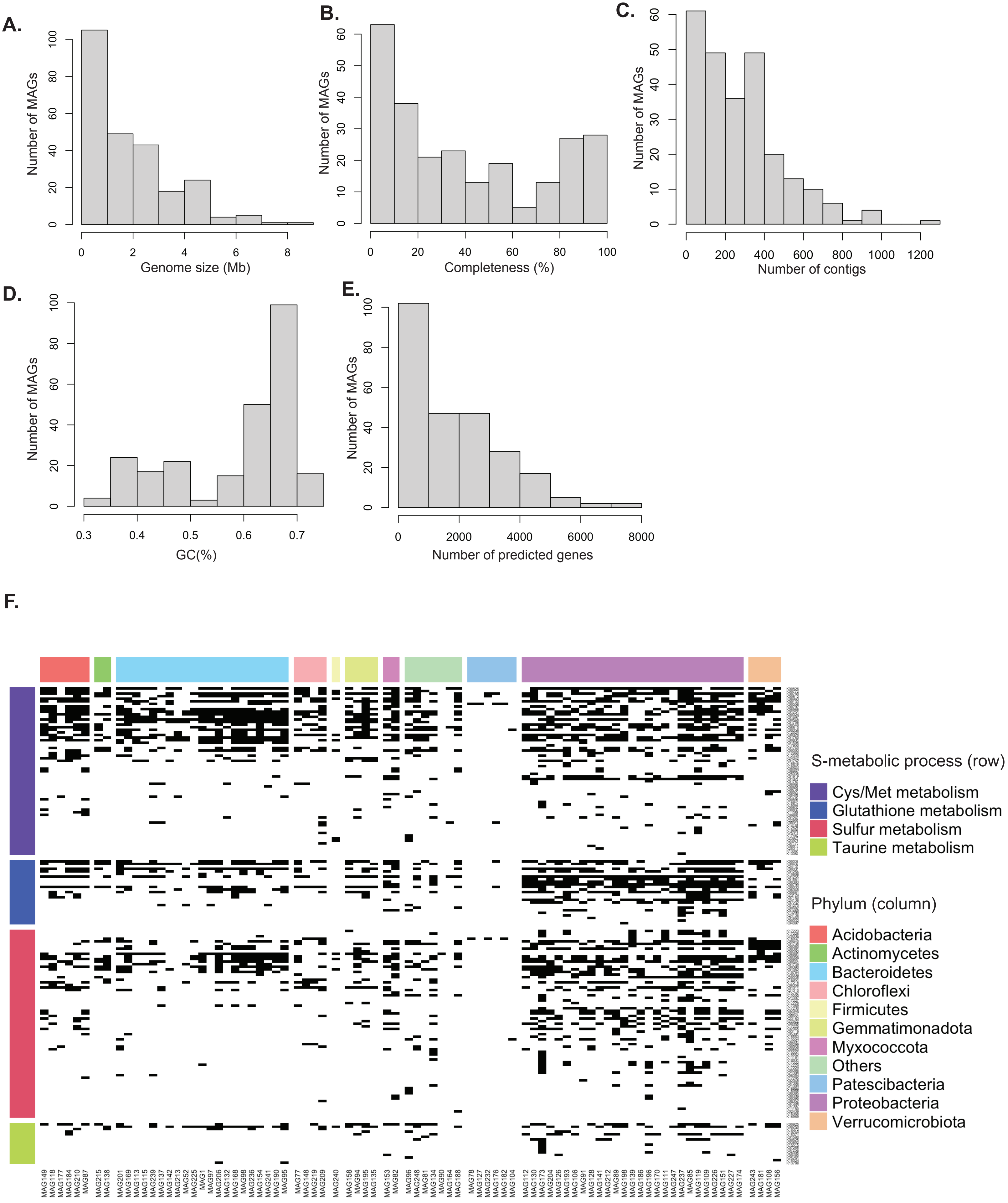
A.
Number of MAGs
Genome size (Mb)
B.
Number of MAGs
Completeness (%)
C.
Number of MAGs
Number of contigs
D.
Number of MAGs
GC(%)
E.
Number of MAGs
Number of predicted genes
F.
S-metabolic process (row)
Cys/Met metabolism
Glutathione metabolism
Sulfur metabolism
Taurine metabolism
Phylum (column)
Acidobacteria
Actinomycetes
Bacteroidetes
Chloroflexi
Firmicutes
Gemmatimonadota
Myxococcota
Others
Patescibacteria
Proteobacteria
Verrucomicrobiota

**Figure S2. Quality profile and S-metabolic potential of metagenome assembled genomes (MAGs) from rhizosphere microbiome of pot-grown plants of selected genotypes.**

Frequency distribution of Genome size (**A**), Completeness (**B**), Number of contigs (**C**), GC content (**D**), and Number of predicted genes (**E**) from the MAGs. **F**. Distribution (presence-absence pattern) of S-metabolic genes (from KEGG) among medium and high-quality MAGs assembled in this study. S-metabolic genes are grouped according to processes (colored groups in rows) and the MAGs are grouped according to their phyla (in columns).

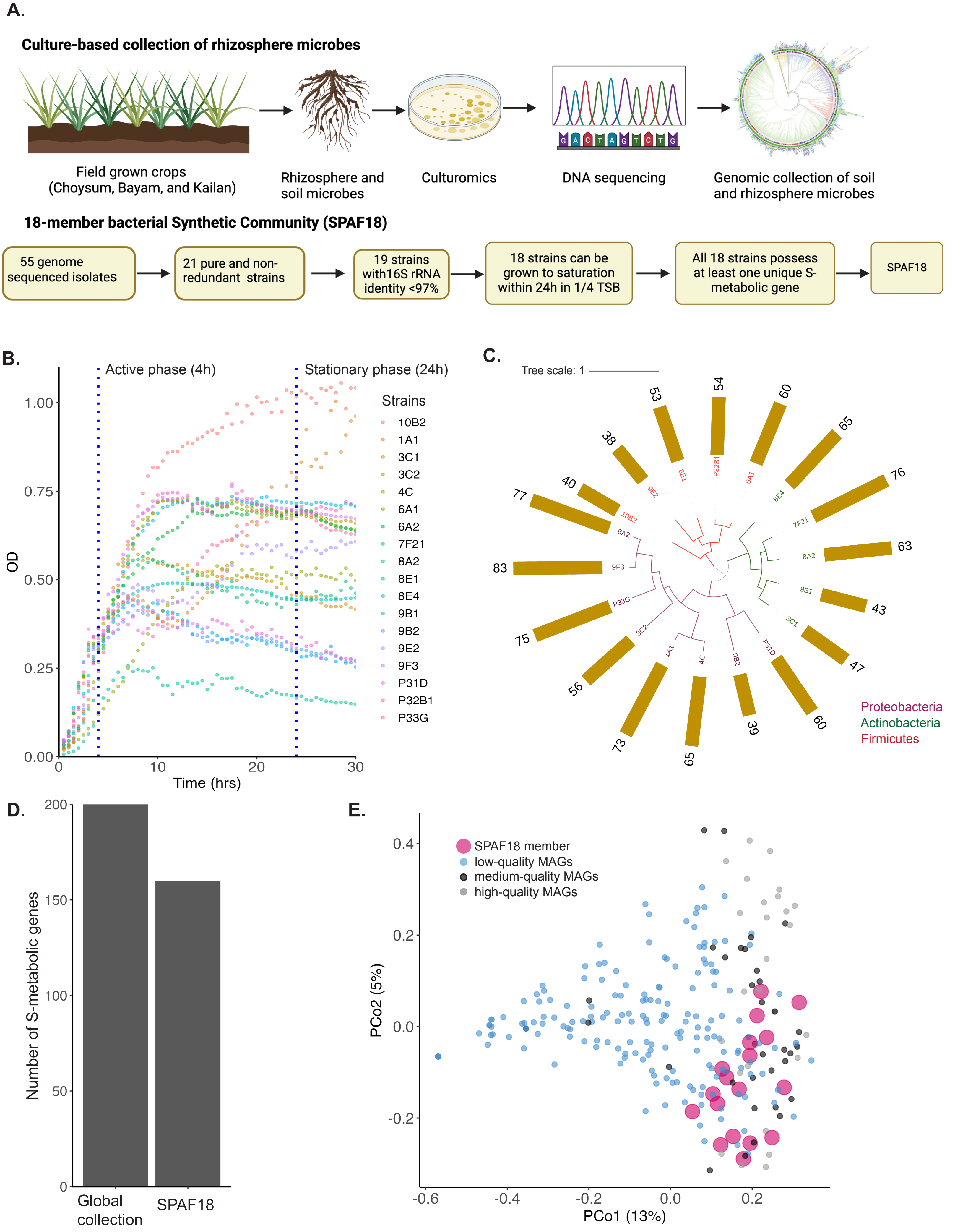
A.
Culture-based collection of rhizosphere microbes
Field grown crops (Choysum, Bayam, and Kailan)
Rhizosphere and soil microbes
Culturomics
DNA sequencing
Genomic collection of soil and rhizosphere microbes
18-member bacterial Synthetic Community (SPAF18)
55 genome sequenced isolates
21 pure and non-redundant strains
19 strains with16S rRNA identity <97%
18 strains can be grown to saturation within 24h in 1/4 TSB
All 18 strains possess at least one unique S-metabolic gene
SPAF18
B.
Active phase (4h)
Stationary phase (24h)
Strains
10B2
1A1
3C1
3C2
4C
6A1
6A2
7F21
8A2
8E1
8E4
9B1
9B2
9E2
9F3
P31D
P32B1
P33G
OD
Time (hrs)
C.
Tree scale: 1
Proteobacteria
Actinobacteria
Firmicutes
D.
Number of S-metabolic genes
Global collection
SPAF18
E.
SPAF18 member
low-quality MAGs
medium-quality MAGs
high-quality MAGs
PCo1 (13%)
PCo2 (5%)

**Figure S3. Workflow for assembling a functionally representative but reduced synthetic rhizosphere bacterial community.**

**A.** Workflow for assembling the 18-membered synthetic rhizosphere bacterial community SPAF18. **B.** Growth rates of individual members of the 18-member synthetic rhizosphere bacterial community SPAF18 in 1⁄4 TSB media. **C.** Phylogenetic tree of SPAF18 members and their total number of S-metabolic genes. **D.** Bar plot showing the total number of S-metabolic genes in SPAF18 and a global collection of rhizosphere bacterial members. **E.** Principal coordinate plot showing the ordination of metagenome assembled genomes (MAGs) and SPAF18 members based on their predicted S-metabolic genes.

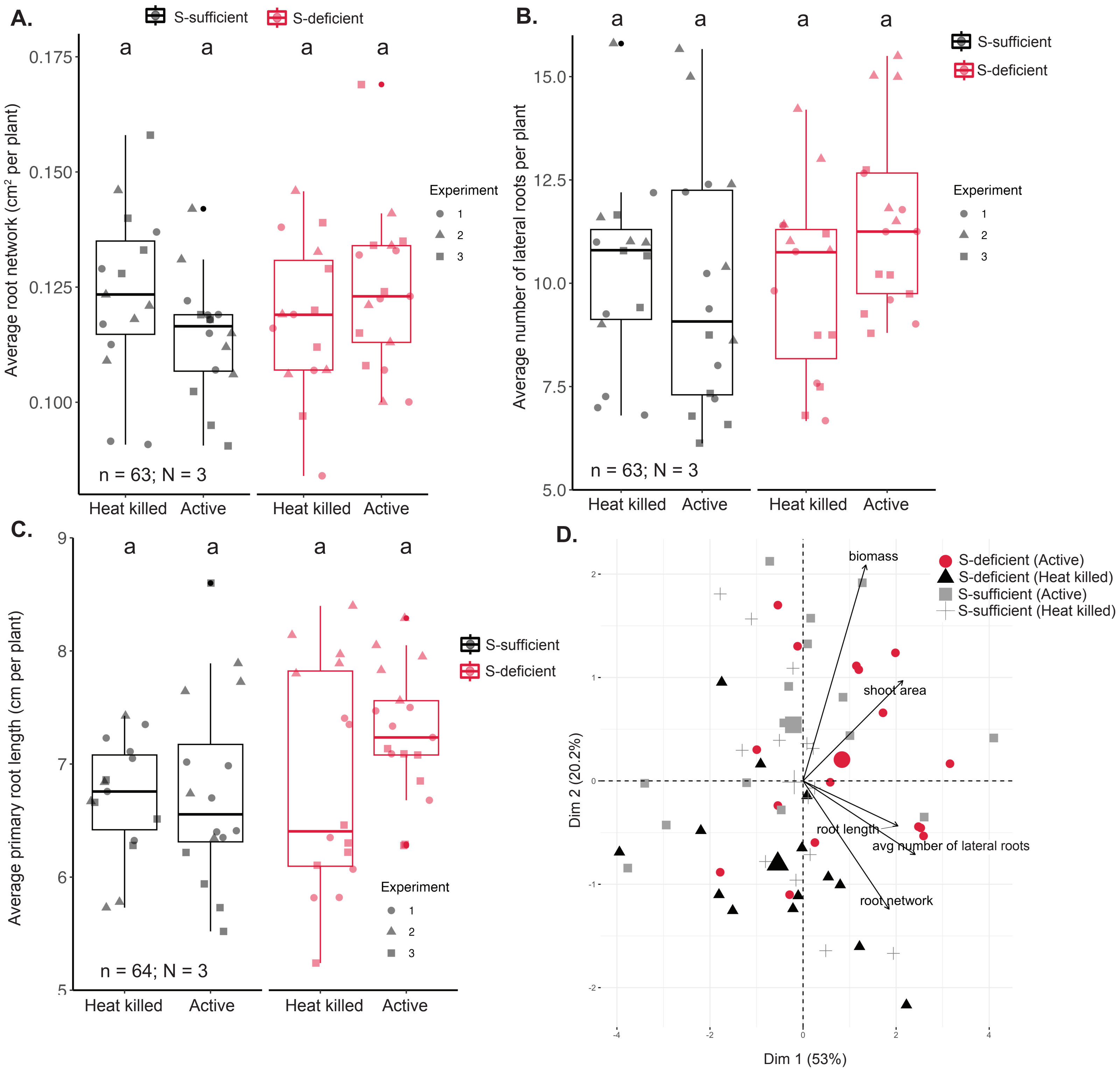
A.
S-sufficient
S-deficient
a
a
a
a
0.175
0.150
0.125
0.100
Average root network (cm² per plant)
Experiment
1
2
3
n = 63; N = 3
Heat killed
Active
Heat killed
Active
B.
a
a
a
a
S-sufficient
S-deficient
15.0
12.5
10.0
7.5
5.0
Average number of lateral roots per plant
Experiment
1
2
3
n = 63; N = 3
Heat killed
Active
Heat killed
Active
C.
9
8
7
6
5
a
a
a
a
S-sufficient
S-deficient
Average primary root length (cm per plant)
Experiment
1
2
3
n = 64; N = 3
Heat killed
Active
Heat killed
Active
D.
biomass
S-deficient (Active)
S-deficient (Heat killed)
S-sufficient (Active)
S-sufficient (Heat killed)
shoot area
root length
avg number of lateral roots
root network
Dim 2 (20.2%)
Dim 1 (53%)
2
1
0
-1
-2
-4
-2
0
2
4

**Figure S4. Effects of SPAF18 on root phenotypes of Arabidopsis plants under S-sufficient and S-deficient conditions.**

**(A)** Average root network ($cm^2$ per plant) **(B)** Average number of lateral roots, and **(C)** Primary root length (cm per plant) of plants inoculated with either heat-killed or active SPAF18 in S- sufficient and S-deficient conditions. Number of biological replicates, n = 63, 63, and 65 for average root network, average number of lateral roots, and average primary root length, respectively, across three independent experiments (N = 3, indicated by shape of data points). **D.** Principal component analysis (PCA) of Arabidopsis phenotypes (fresh biomass, shoot area, primary root length, root network, and average number of lateral roots) based on samples treated with either heat killed or active SPAF18 under both S-sufficient and S-deficient conditions. Arrows indicate the loadings of the phenotypes separating the samples on both PCs (Dim 1 and Dim 2). PCA is performed by pooling data from three independent experiments (N = 3).

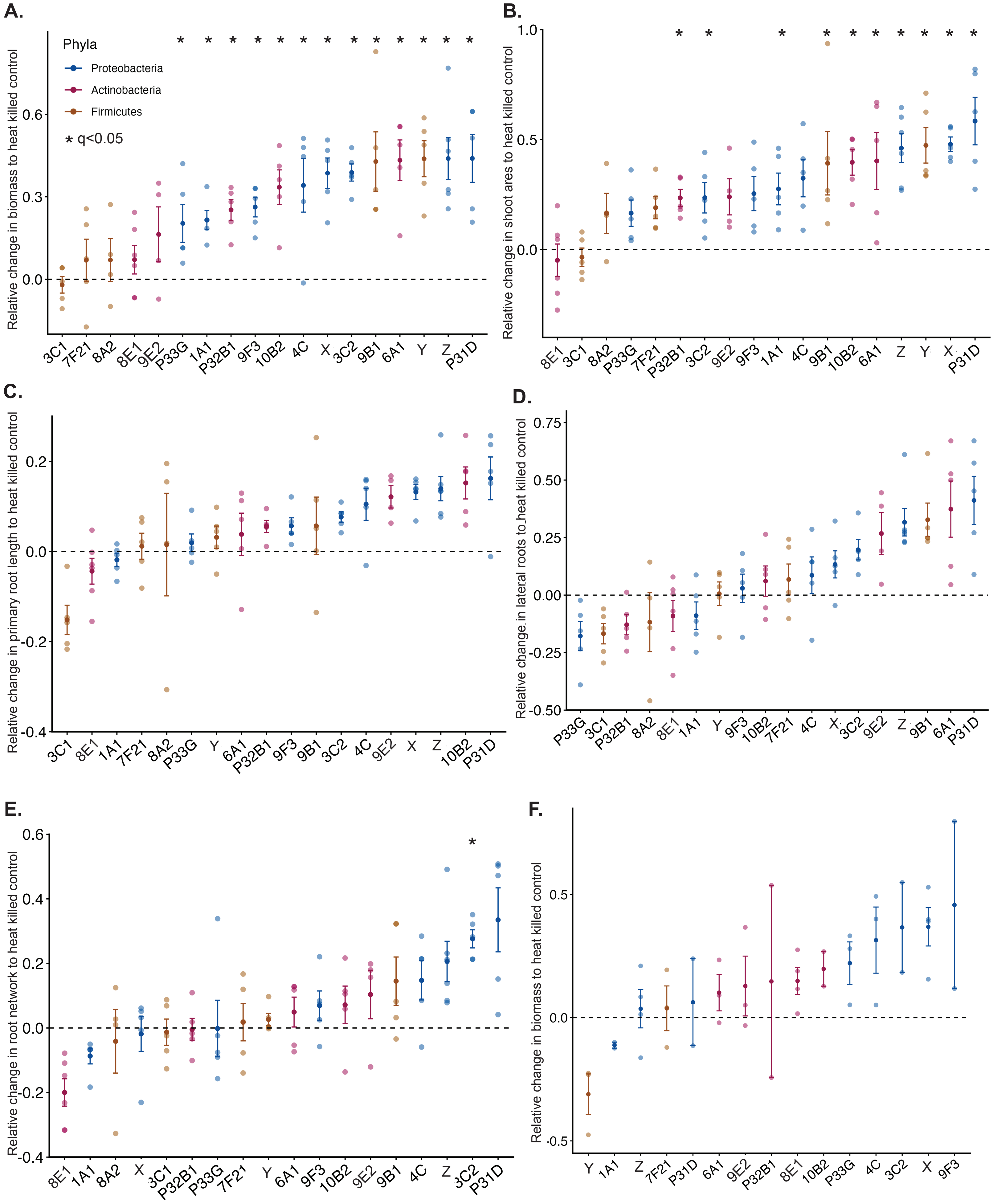

A.
Phyla
Proteobacteria
Actinobacteria
Firmicutes
* q<0.05
Relative change in biomass to heat killed control
3C1 7F21 8A2 8E1 9E2 P33G 1A1 P32B1 9F3 10B2 4C X 3C2 9B1 6A1 Y Z P31D
B.
Relative change in shoot ares to heat killed control
8E1 3C1 8A2 P33G 7F21 P32B1 3C2 9E2 9F3 1A1 4C 9B1 10B2 6A1 Z Y X P31D
C.
Relative change in primary root length to heat killed control
3C1 8E1 1A1 7F21 8A2 P33G Y 6A1 P32B1 9F3 9B1 3C2 4C 9E2 X Z 10B2 P31D
D.
Relative change in lateral roots to heat killed control
P33G 3C1 P32B1 8A2 8E1 1A1 Y 9F3 10B2 7F21 4C X 3C2 9E2 Z 9B1 6A1 P31D
E.
Relative change in root network to heat killed control
8E1 1A1 8A2 X 3C1 P32B1 P33G 7F21 Y 6A1 9F3 10B2 9E2 9B1 4C Z 3C2 P31D
F.
Relative change in biomass to heat killed control
Y 1A1 Z 7F21 P31D 6A1 9E2 P32B1 8E1 10B2 P33G 4C 3C2 X 9F3

**Figure S5. Effects of individual members of SPAF18 on all five Arabidopsis phenotypes measured under S-deficiency.**

Relative increase in fresh biomass **(A)**, shoot area **(B)**, primary root length **(C),** root network **(D)**, and number of lateral roots (**E)** for individual members of SPAF18 relative to heat killed control. Boxes and data points are colored according to phyla. Statistical significance is calculated based on ANOVA followed by post-hoc Tukey's test (FDR corrected p-value or $q < 0.05$). **F.** Relative increase in fresh biomass for individual members of SPAF18 under S-sufficient condition (3C1, 9B1, and 8A2 not tested). Statistical significance is calculated same as in A-E.

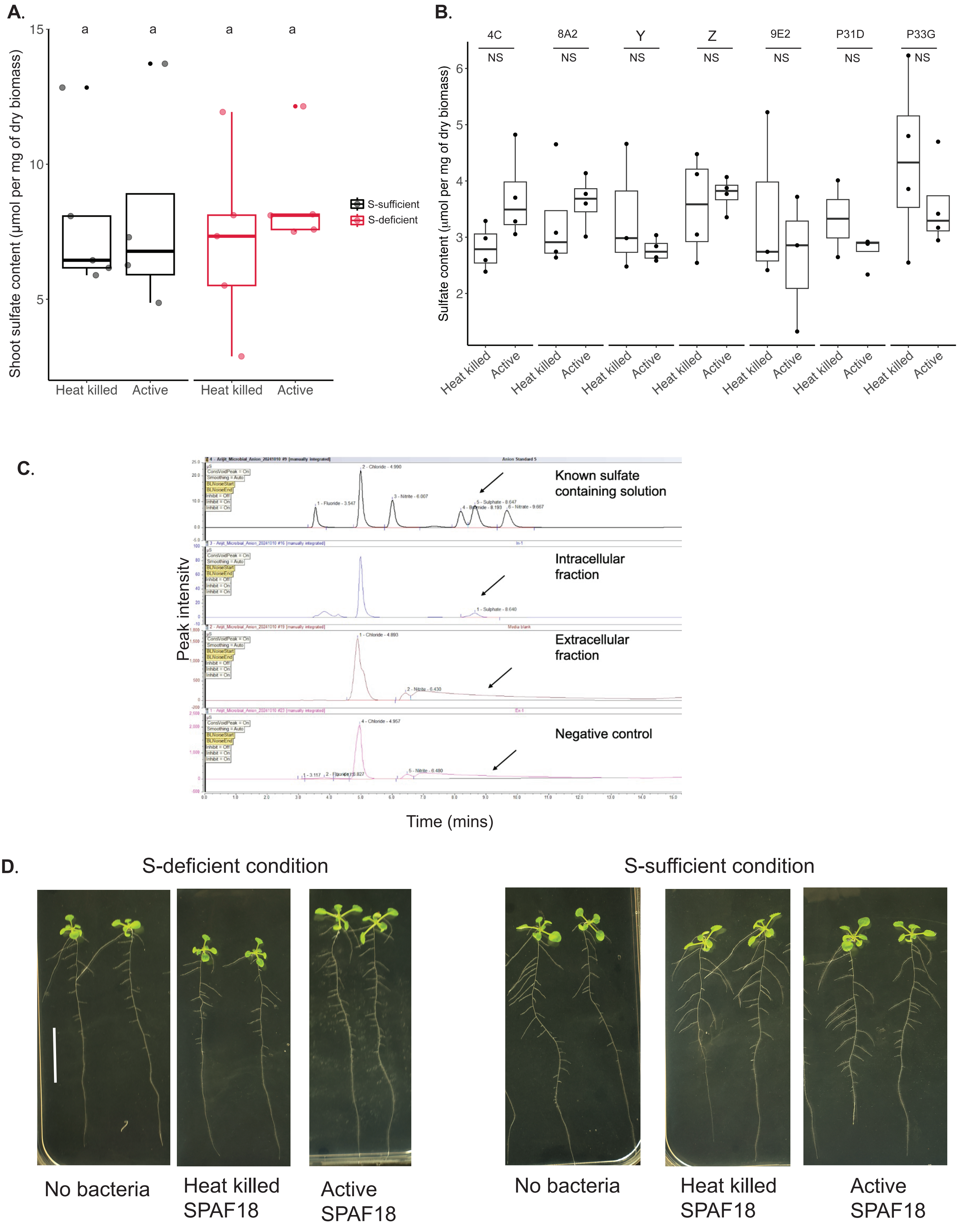
A.
Shoot sulfate content (µmol per mg of dry biomass)
15
10
5
a
a
a
a
S-sufficient
S-deficient
Heat killed
Active
Heat killed
Active
B.
Sulfate content (µmol per mg of dry biomass)
6
5
4
3
2
4C
8A2
Y
Z
9E2
P31D
P33G
NS
NS
NS
NS
NS
NS
NS
Heat killed
Active
Heat killed
Active
Heat killed
Active
Heat killed
Active
Heat killed
Active
Heat killed
Active
Heat killed
Active
C.
Peak intensity
Known sulfate containing solution
Intracellular fraction
Extracellular fraction
Negative control
Time (mins)
D.
S-deficient condition
S-sufficient condition
No bacteria
Heat killed SPAF18
Active SPAF18
No bacteria
Heat killed SPAF18
Active SPAF18

**Figure S6. Sulfate content of Arabidopsis plants treated with SPAF18 or its individual members.**

**A.**Sulfate content of 12-days old Arabidopsis seedlings treated with either heat killed or active SPAF18. Boxes and data points are colored according to the sulfate levels. **B.** Sulfate content of 12-days old Arabidopsis seedlings treated with selected individual members compared to their heat killed control. Number of biological replicates (n = 2 to 4) per strain. **C.** Raw chromatogram (from ion-chromatography) showing sulfate levels in the extracellular and intracellular fractions of SPAF18. Sulfate is not detected in negative controls and extracellular fraction of SPAF18. **D.** Representative images of 12-days old Arabidopsis plants treated with no-bacteria, heat killed, and active SPAF18 in both S-sufficient and S-deficient conditions. Scale bar represents 2 cm length.

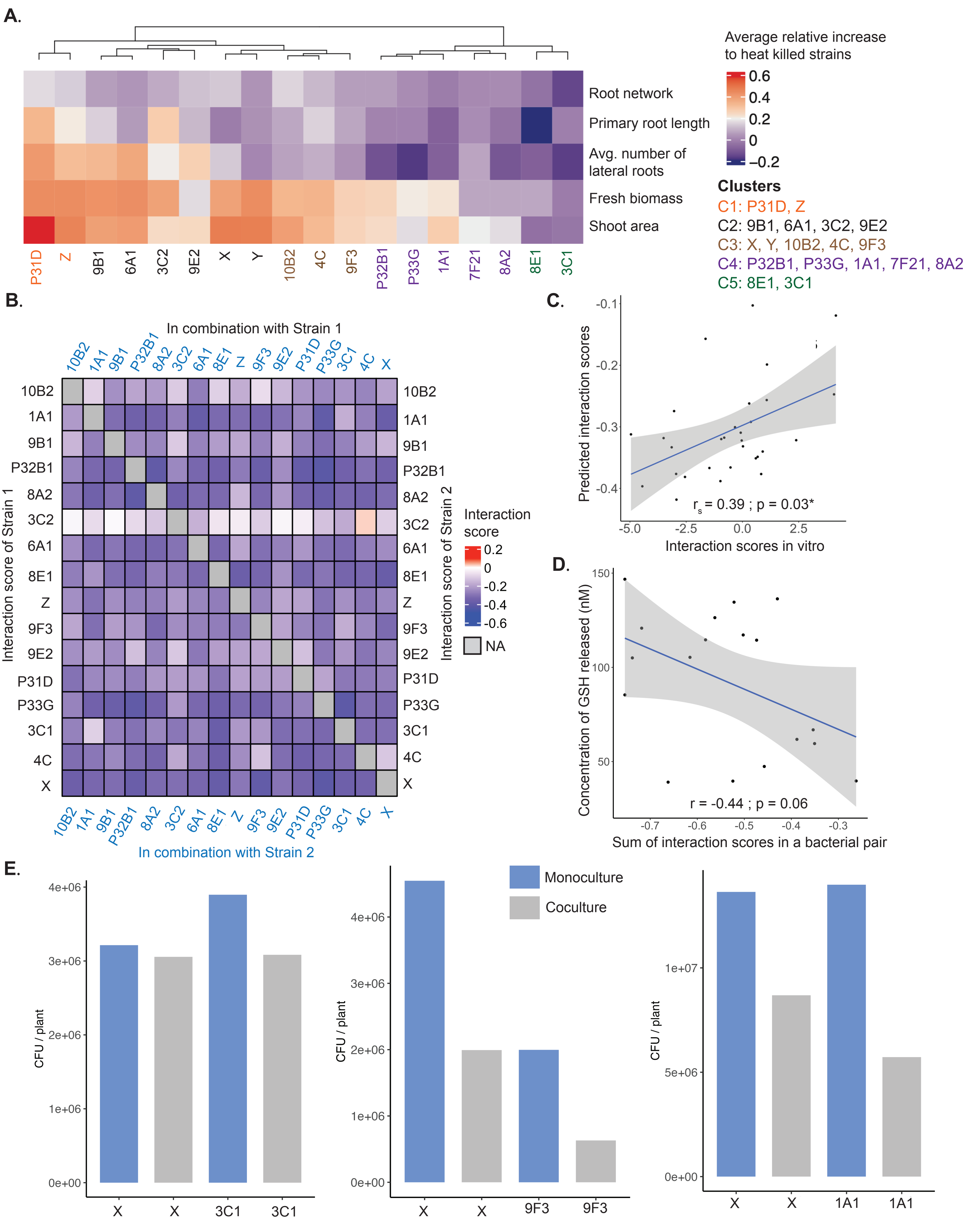
A.
Average relative increase to heat killed strains
0.6
0.4
0.2
0
−0.2
Root network
Primary root length
Avg. number of lateral roots
Fresh biomass
Shoot area
P31D
Z
9B1
6A1
3C2
9E2
X
Y
10B2
4C
9F3
P32B1
P33G
1A1
7F21
8A2
8E1
3C1
Clusters
C1: P31D, Z
C2: 9B1, 6A1, 3C2, 9E2
C3: X, Y, 10B2, 4C, 9F3
C4: P32B1, P33G, 1A1, 7F21, 8A2
C5: 8E1, 3C1
B.
In combination with Strain 1
10B2
1A1
9B1
P32B1
8A2
3C2
6A1
8E1
Z
9F3
9E2
P31D
P33G
3C1
4C
X
Interaction score of Strain 1
Interaction score of Strain 2
Interaction score
0.2
0
-0.2
-0.4
-0.6
NA
In combination with Strain 2
C.
Predicted interaction scores
-0.1
-0.2
-0.3
-0.4
$r_s$ = 0.39 ; p = 0.03*
-5.0
-2.5
0.0
2.5
Interaction scores in vitro
D.
Concentration of GSH released (nM)
150
100
50
r = -0.44 ; p = 0.06
-0.7
-0.6
-0.5
-0.4
-0.3
Sum of interaction scores in a bacterial pair
E.
CFU / plant
4e+06
3e+06
2e+06
1e+06
0e+00
X
X
3C1
3C1
Monoculture
Coculture
X
X
9F3
9F3
1e+07
5e+06
X
X
1A1
1A1

**Figure S7. Clustering of SPAF18 members for machine learning models, their pairwise interactions, sulfate content, and Arabidopsis phenotypes treated with SPAF18**

**A.** Hierarchical clustering of SPAF18 members based on their effects on plant phenotypes under S-deficiency. The colors within the heatmap represents the scaled relative increase to heat killed control for each strain. Strains are colored according to the cluster they were assigned. These clusters were considered for stratifying the bacterial members into modules while developing machine learning models. **B.** Heatmap showing interaction scores among selected pairs of SPAF18. Colors indicate the interaction scores predicted from their simulated growth rates in monoculture and coculture (described in Figure 5A). For a bacterial pair, such as for 10B2 and 1A1, the interaction score for 1A1 is indicated in the first element of the second row in the heatmap. Whereas the interaction score of 10B2 is indicated on the second element of the first row. **C.** Correlation between predicted (from genome-scale metabolic models; Table S13) and in vitro experimental (strains grown in Artificial root exudates + MS media; Table S14) interaction scores. Data points are shown for individual interaction scores in a bacterial pair. **D.** Correlation between the concentration of GSH released in the extracellular space of selected bacterial pairs in coculture (Table S17) and their interaction scores. Each data point represents the median GSH concentration from n = 3 biological replicates. **E.** Bar plot showing the colony forming units (CFUs) per Arabidopsis roots in monoculture and coculture of selected bacterial pairs. Average number of CFUs are shown (n = 2/3). Detailed dataset is presented in Table S15.
